# Thymic self-recognition-mediated TCR signal strength modulates antigen-specific CD8^+^ T cell pathogenicity in non-obese diabetic mice

**DOI:** 10.1101/2024.06.10.596762

**Authors:** Chia-Lo Ho, Li-Tzu Yeh, Yu-Wen Liu, Jia-Ling Dong, Huey-Kang Sytwu

## Abstract

Our understanding of autoimmune diabetes underscores the critical involvement of CD8^+^ T cells recognizing islet-specific antigens. However, the influence of thymic positive selection on diabetogenic CD8^+^ T cell development remains unclear. Using CD5 marker representing T-cell receptor (TCR) signal strength, we illustrated that naïve CD5^hi^CD8^+^ T cells of non-obese diabetic (NOD) mice with enhanced TCR signals displayed predisposed differentiated/memory T cell traits with increased activation and proliferation upon TCR stimulation, compared to CD5^lo^ counterparts. Additionally, CD5^hi^CD8^+^ T cells exhibited gene expression landscape similar to effector T cells and exacerbated disease in transfer model. Interestingly, the protective effects of transgenic phosphatase Pep expression, which lowers TCR signaling and diabetes incidence, were abolished in NOD strain 8.3 with high CD5 expression linked to increased thymic positive selection. Strikingly, TCR repertoire analysis identified higher frequencies of autoimmune disease-related clonotypes in naïve CD5^hi^CD8^+^ cells, supporting that distinct effector functions arise from intrinsic TCR repertoire differences. Overall, CD5^hi^CD8^+^ clones may be potential targets for autoimmune diabetes treatment.

## Introduction

T cells play a crucial role in inducing immunity by recognizing peptides presented on MHC molecules through their TCR during pathogen infection. The strength of the TCR’s reactivity to the foreign antigen is contributed by both the affinity of the TCR for that antigen and the basal TCR signaling level of the T cell, which is determined by weak interactions between the TCR and self-peptides presented during T cell development (1, 2). Previous studies have shown that T cell self-reactivity with weak interaction between TCR and self-peptide-MHC (self-pMHC) not only instructs T cell development and activation thresholds, but also influences their functional response (2–4). It is noteworthy that the strength of the weak interaction can be evaluated through CD5, whose expression level is considered proportional to T cell self-reactivity in the developmental stage, and may reflect the responses of T cells to foreign-pMHC complexes in the future (2, 5, 6). CD5 is a scavenger receptor cysteine-rich protein expressed on T and B cells that regulates

TCR and B-cell receptor signaling (7, 8). It functions as a scaffold protein for the recruitment of phosphatase SHP-1 and E3-ubiquitin ligases CBL and CBLB, to reduce the phosphorylation of proximal TCR signaling molecule Zap70, thereby inhibiting TCR-mediated signaling (9). CD5, as a negative regulator for TCR signaling, also plays a crucial role in fine-tuning the process of positive and negative selection during thymocyte development (8, 10). Specifically, thymocytes with higher self-reactivity TCRs induce CD5 expression, which helps them adapt to strong TCR signaling and reduces the likelihood of negative selection. This, in turn, promotes thymocyte positive selection and broadens the affinity spectrum of the TCR repertoire during the TCR self- recognition process. Therefore, the expression of CD5 is not only a surrogate marker for the strength of TCR-self-peptide-MHC interactions during development but also reflects the intrinsic response set point of T cells to a signal (5, 10). This selection process enriches the mature CD4^+^ T cell repertoire with clones that exhibit increased responses to self-peptide-MHC, thereby displaying enhanced peripheral immune responses to pathogens and autoantigens *in vivo* (5, 11, 12). Not only in CD4^+^ T cells, previous study has also shown that CD5^hi^ population in CD8^+^ T cells, characterized by higher reactivity to self-peptide-MHC, are prone to positive selection (5, 13). In diabetes-prone NOD mice, though negative selection in central tolerance limits the early contribution of high-avidity T cells to disease development, prolonged inflammation in pancreatic islets still drives the avidity maturation of islet-specific glucose-6-phosphatase catalytic subunit-related protein (IGRP)-specific CD8^+^ T cells, leading to increased cytotoxicity towards β cells (14–16). This suggests that mechanisms other than incomplete thymic negative selection are involved in generating autoreactive T cells. Here, we investigate whether the CD5-correlated self-pMHC reactivity, which attributes to TCR signal strength in autoreactive CD8^+^ T cells during thymic positive selection, contributes to the development of autoimmune disease.

The NOD mouse model is commonly used for studying autoimmune diabetes. NOD mice spontaneously develop autoimmune diabetes similar to human type 1 diabetes (T1D) with immune-mediated destruction of β cells in the pancreas (17–20). Additionally, genetic variation influences cell lineage decisions during T cell development, affecting positive and negative selection processes and contributing to the emergence of pathogenic CD4^+^ T cells in diabetes-prone NOD mice (16). Both CD4^+^ and CD8^+^ T cells play critical roles in pathogenic diabetogenesis, in which CD4^+^ T cells drive disease onset and CD8^+^ T cells mediate β cell destruction (21). However, little is known about the factors affecting CD8^+^ T cell activity, and the extent to which CD8^+^ T cell reactivity to self-pMHC during thymic positive selection affects its responses to autoantigens during autoimmune processes remains unclear.

We hypothesize that the spectrum of CD8^+^ T cell diabetogenicity correlates with its expression level of CD5, representing the magnitude of self-pMHC-mediated TCR signaling. In this study, we use *Ptpn22* (22–24) and islet antigen-specific 8.3 TCR transgenic (NOD8.3) (25, 26) mice in addition to conventional NOD mice to investigate this hypothesis. The *Ptpn22* gene encodes PEST domain-enriched tyrosine phosphatase (Pep), which negatively regulates TCR signaling. We have previously demonstrated that different modulation of Pep in effector and regulatory CD4^+^ T cells leads to attenuation of autoimmune diabetes in distal Lck promoter-driven Pep transgenic NOD mice (dLPC/NOD) (27). Here, we crossed dLPC/NOD to NOD8.3 transgenic mice to generate dLPC/NOD8.3 double transgenic mice and further investigate whether transgenic Pep-mediated protective potential in CD4^+^ T cells with lower TCR signaling (27) can also be applied to an autoantigen-specific CD8^+^ T cells with higher CD5 and TCR basal signaling in 8.3 TCR transgenic mice. Our data reveal that the activation and proliferation of CD8^+^ T cells from dLPC/NOD8.3 mice was not attenuated, compared to CD8^+^ T cells from NOD8.3 mice, suggesting that a dominant role of autoantigen reactivity in CD8^+^ T cell with high TCR basal signaling and CD5 overrides the transgenic Pep-mediated signaling attenuation during autoimmune diabetogenesis. Exploring a potential modulation of CD5-dependent TCR basal signaling and its impact on T cell reactivity may offer a promising therapeutic strategy for the treatment of T1D in the future.

## Results

### Phenotypic difference exists in naïve CD5^hi^CD8^+^ and CD5 ^lo^CD8^+^ T cells of NOD mice

Given that CD5 can serve as a surrogate marker of TCR basal signal strength, which correlates with T cell’s self-recognition capability, our initial analysis aimed to determine whether naïve (CD44^lo^CD62L^hi^) CD8^+^ T cells from NOD mice with different CD5 levels (CD5^hi^CD8^+^ and CD5^lo^CD8^+^ T cells) display distinct levels of the TCR basal signal markers p-CD3ζ and p-Erk. The histograms and gMFI analysis revealed significantly increased levels of p-CD3ζ and p-Erk in CD5^hi^CD8^+^ T cells, in comparison to CD5^lo^CD8^+^, while the amounts of p-CD3ζ and p-Erk in total naive CD8^+^ T cells fell between the CD5^hi^ and CD5^lo^ subsets (Figure 1, A and B, and Supplementary Figure 1A) in both spleen and pancreatic lymph nodes (PLNs). This result implies an enhanced signaling potential and activation status of CD5^hi^CD8^+^ T cells, compared to CD5^lo^CD8^+^ cells, in NOD mice. Since two critical T-box transcription factors, T-bet and Eomes, are regulators of differentiation and effector functions of CD8^+^ T cells (28–30), we further investigated whether intrinsic differences in self-recognition capabilities and subsequent functional heterogeneity between CD5^hi^CD8^+^ and CD5^lo^CD8^+^ populations correlate the expression levels of T-bet and Eomes. Our results indicated higher levels of T-bet and Eomes in the naïve CD5^hi^CD8^+^ population in both spleen and PLNs, compared to CD5^lo^ counterparts (Figure 1C and Supplementary Figure 1B). Elevated expressions of T-bet and Eomes in CD8^+^ T cells have been reported to play critical role in promoting T cell migration to inflamed tissues through induction of chemokine receptors (31, 32) in addition to their association with the differentiation of activated CD8^+^ T cells (28). These upregulated T-bet and Eomes in naïve CD5^hi^CD8^+^ T cells (Figure 1C) were consistent with elevated levels of Granzyme B, TNF-α, IFN-γ and IL-2 (Figure 1D and Supplementary Figure 1C), indicating the potential of these cells in cytokine production and effector functions. Additionally, CD5^hi^CD8^+^ subset expressed higher level of chemokine receptor CXCR3 (Figure 1E) than CD5^lo^CD8^+^ counterparts, a characteristic often associated with effector and memory T cell populations, implying a enhanced ability for migration and localization in response to specific chemotactic signals.

**Figure 1.**
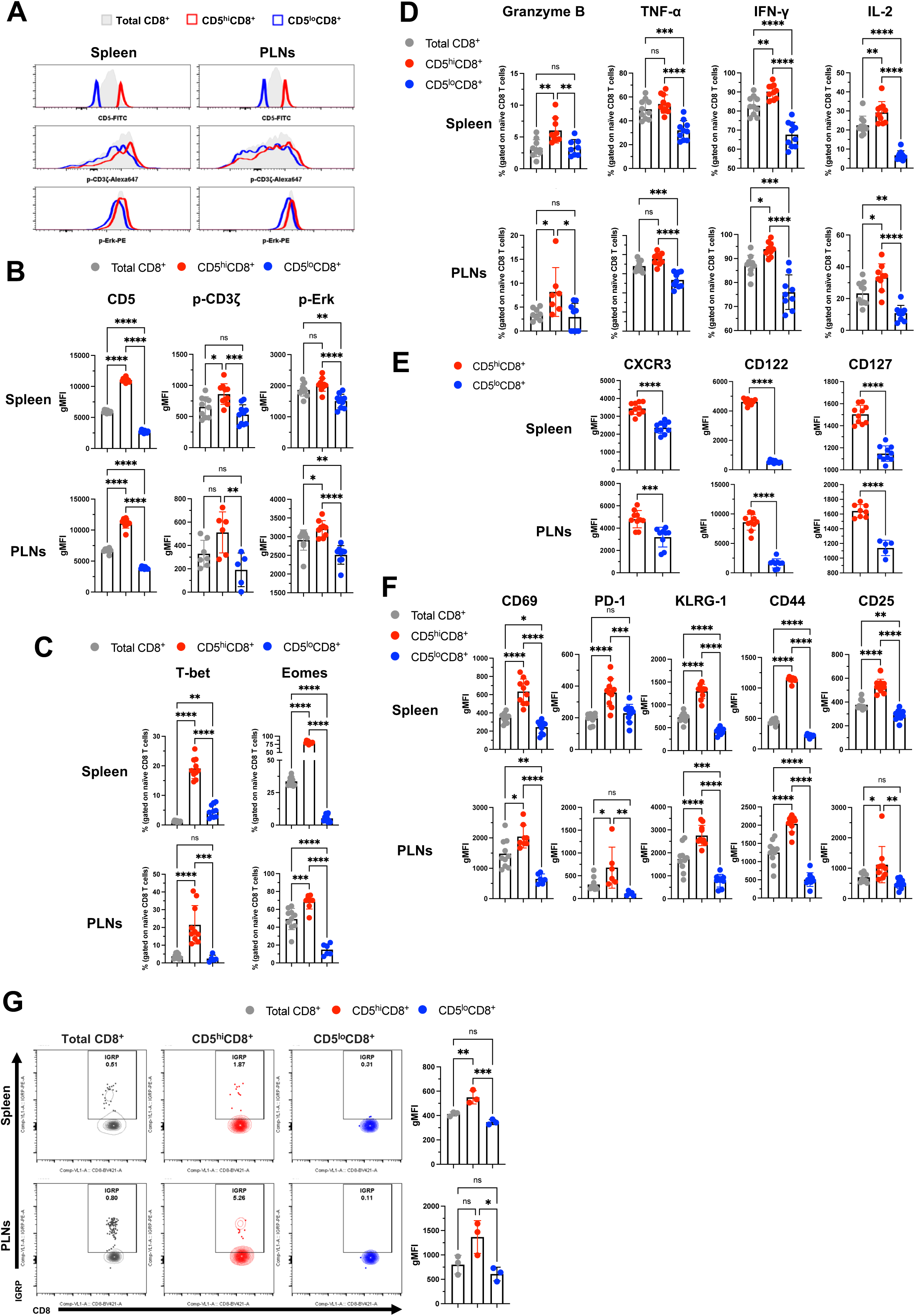
Unique phenotypic traits of naïve NOD CD8^+^ T cells are stratified by CD5 expression. In (**A**) to (**G**), flow cytometry analysis of total naïve CD8^+^ T cells (CD44^lo^CD62L^hi^; grey), and the upper 5% (CD5^hi^CD8^+^; red) and lower 5% (CD5^lo^CD8^+^; blue) based on CD5 expression, from the spleen and PLNs of normoglycemic female NOD mice aged 6 to 8 weeks. (**A**) Representative flow cytometry histograms and the corresponding (**B**) geometric mean fluorescence intensity (gMFI) of CD5, p-CD3ζ and p-Erk expression shown in the indicated samples. Additionally, the percentages of (**C**) transcription factors T-bet and Eomes, along with (**D**) cytokines Granzyme B, TNF-α, IFN-γ and IL-2 in CD8^+^ T cells are also presented in the indicated samples. The gMFI for effector/memory T cell-related markers includes (**E**) CXCR3, CD122, CD127 and (**F**) CD69, PD-1, KLRG-1, CD44 and CD25. (**G**) Representative flow cytometry plots (left) and gMFI (right) of positive IGRP-tetramer staining in CD8^+^ T cells compared among naive total CD8^+^ T cells, CD5^hi^CD8^+^ and CD5^lo^CD8^+^ populations. Data represent mean ± SD. The data presented in (**A**) to (**F**) represent two experiments. The sample size was n = 5-10 (**B**), n = 6-10 (**C**), n = 7-10 (**D**), n = 5-10 (**E**) n = 6-10 (**F**) and n = 3 (**G**) per group. \**P* < 0.05, \*\**P* < 0.01, \*\*\**P* < 0.001, \*\*\*\**P* < 0.0001, by one-way ANOVA (**B**–**D**, **F**, **G**) and unpaired, two-tailed t test (**E**).

Building upon our observations of the phenotypic and functional differences between naïve CD5^hi^ and CD5^lo^CD8^+^ T cell subsets in NOD mice (Figure 1, A-D), as well as previous research highlighting T-bet- and Eomes-related regulatory mechanisms within CD8^+^ T cell populations (3, 28), we next sought to investigate whether CD5-associated self-recognition capabilities influence specific gene regulation in CD8^+^ T cells via distinct levels of T-bet and Eomes. It is well documented that T-bet and Eomes work together to control CD122 expression, a common β chain receptor for IL-2 and IL-15, which is crucial for the survival of IL-15-dependent memory CD8^+^ T cells (30, 33). Additionally, CD122, in cooperation with CD127 (IL-7Rα), represents essential receptors for IL-2, IL-15, and IL-7. They play a pivotal role in regulating CD8^+^ T cell subsets during the memory phase of the immune response (34–37), which is crucial for maintaining memory T cell homeostasis. As shown in Figure 1E, higher expression of CD122 and CD127 was observed in the CD5^hi^CD8^+^ population, supporting that the overall phenotypic traits of naïve CD5^hi^CD8^+^ T cells resemble those of memory CD8^+^ T cells (Figure 1E), particularly in their ability to maintain T cell homeostasis.

In addition to distinct expression patterns of these memory T cell-related homeostasis markers, like CD122 and CD127 (Figure 1E), we also identified higher expressions of other T cell activation markers, including CD69 (38, 39), PD-1 (40, 41), and KLRG-1 (37, 42), in naïve CD5^hi^CD8^+^ T cells compared to their CD5^lo^ counterparts (Figure 1F). Similar to PD-1, the activation marker CD44, an adhesion molecule and marker for antigen-experienced T cells, was up-regulated in the CD5^hi^CD8^+^ population (Figure 1F), as previously reported (3, 34, 43, 44). Additionally, CD5^hi^ cells also exhibit elevated expression of CD25 (IL-2Rα), supporting the presence of intrinsic phenotypic distinction between the CD5^hi^ and CD5^lo^ populations in NOD naïve CD8^+^ T cells (Figure 1F). Since the increased number of islet antigen IGRP-specific CD8^+^ T cells positively correlate with insulitis progression both in human T1D and NOD mice, we thus used MHC-tetramer technique to trace IGRP_206–214_-specific CD8^+^ T cells in NOD mice. Our results revealed that CD5^hi^CD8^+^ T cells compared to CD5^lo^CD8^+^ counterpart had significant increase in IGPR-tetramer staining level (Figure 1G and Supplementary Figure 1D), suggesting that CD5^hi^CD8^+^ T cells with higher self-antigen recognizing ability contribute more to islet-destructive pathogenesis in T1D. To further evaluate antigen specificity, we also examined Insulin B15-23 (InsB15–23)-specific CD8^+^ T cells using MHC-pentamer staining and observed increased frequencies in the CD5^hi^ subset compared to CD5^lo^ and total naïve CD8^+^ T cells (Supplementary Figure 1E), reinforcing their enriched autoreactive potential. Collectively, in line with previous findings, our results indicate that the phenotypic characteristics of naïve CD5^hi^CD8^+^ T cells resemble those of effector/memory T cells and confer distinct functional capabilities in autoimmune diabetogenesis compared to CD5^lo^CD8^+^ cells.

### Differential gene expression profiling of naïve CD8^+^ T cells stratified by CD5 levels identifies autoimmune disease-associated gene signatures

To unveil the underlying differences in phenotypic characteristics between CD5^hi^CD8^+^ and CD5^lo^CD8^+^ T cells, we analyzed the gene expression profiling by sorting the 10% upper and lower CD5-expressing naïve CD8^+^ T cells from prediabetic mice at the age of 6 to 8 weeks old (Figure 2A). We performed differentially expressed genes (DEGs) analysis using multiple platforms and bioinformatic tools to compare the transcriptomic profiles of CD5^hi^ versus CD5^lo^ T cells. Our DEGs results showed that a total of 185 genes were found to be differentially expressed between CD5^hi^ and CD5^lo^, with a fold change greater than 2 and a p-value less than 0.05 cutoffs; within these genes, 133 were upregulated and 52 were downregulated (Supplementary Table 1 and 2). The significantly upregulated DEGs revealed by the volcano plot (Figure 2B) in CD5^hi^ subset include memory T cell homeostasis marker *Il2rb* (CD122), T cell effector/memory function-associated markers *Tbx21*, *Eomes*, and chemokine receptor *Cxcr3*, consistent with the results obtained in Figure 1. Interestingly, the *Xcl1* gene, which encodes the ligand XCL1 exclusively binding to XCR1 on dendritic cells and facilitating antigen-specific CD8^+^ T cell expansion, IFN- γ secretion and cytotoxicity, was also upregulated in the CD5^hi^ population (28, 45, 46), implying that CD5^hi^CD8^+^ T cells are poised for efficient priming upon antigen encounter.

**Figure 2.**
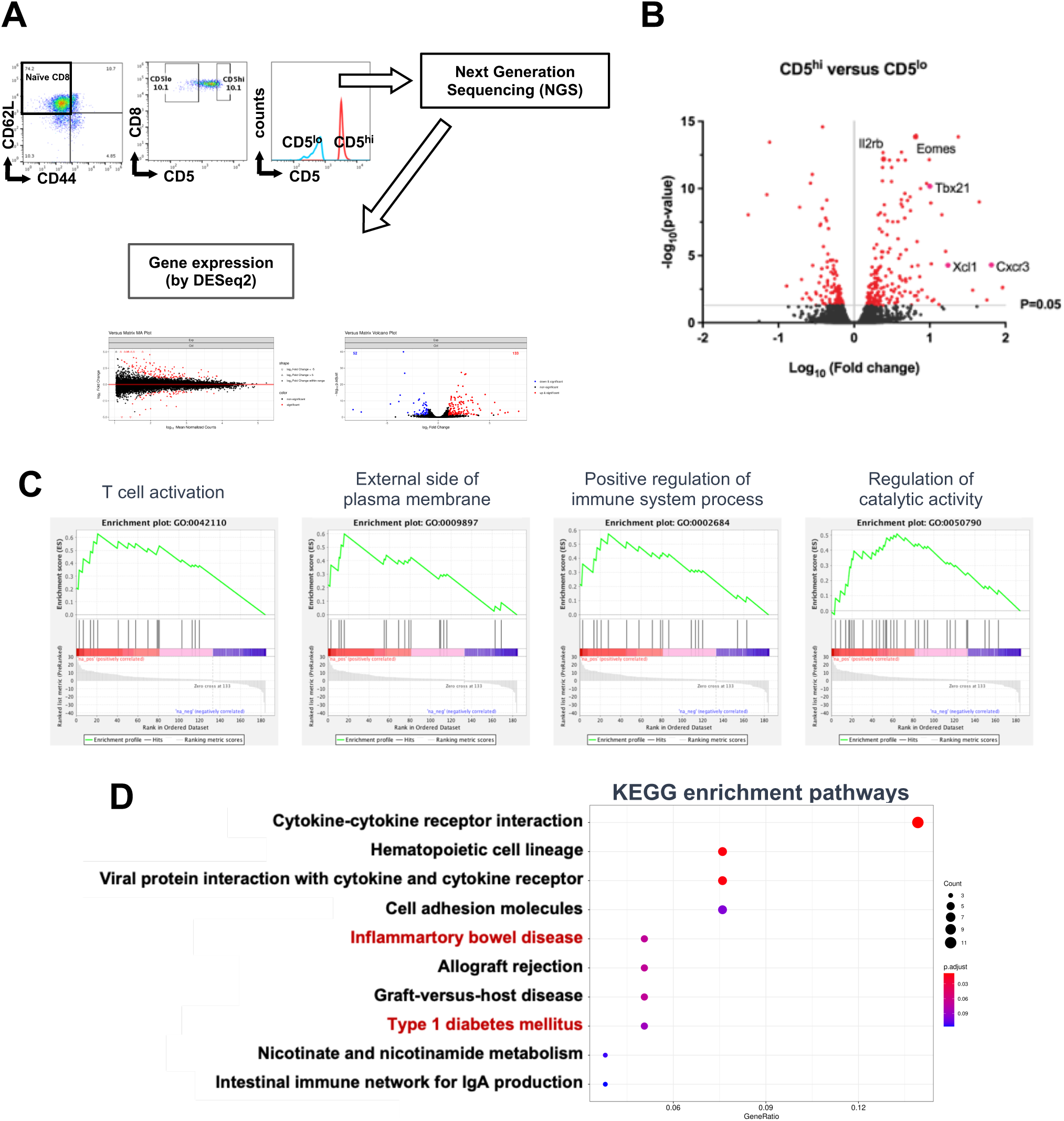
The gene expression profile of naïve CD5^hi^CD8^+^ T cells reveals a poised phenotype for effector T cells in autoimmune diseases. (**A**) RNA-Seq performed on the sorted 10% upper and lower CD5-expressing of naïve CD8^+^ T cells (CD5^hi^CD8^+^ and CD5^lo^CD8^+^) from 6-8-week-old normoglycemia female NOD mice to understand the gene expression profile of CD5^hi^CD8^+^ T cells and comparing that to the CD5^lo^ counterpart to gain insights into their potential role in autoimmune diseases. (**B**) The volcano plot displaying the transcripts that are upregulated and downregulated by RNA sequencing of naïve CD5^hi^CD8^+^ T cells compared to CD5^lo^CD8^+^ T cells. The plot shows the log10 fold change of each transcript and the significance of the change, measured by the negative log10 of the adjusted *P*-value (adj.p). Genes with higher fold changes and lower adj.p values (red plots) indicate a significant change in expression between the two groups. (**C**) The gene set enrichment score plot for gene ontology terms (T cell activation, external side of plasma membrane, positive regulation of immune system process, and regulation of catalytic activity). Higher scores indicate a higher degree of enrichment of these processes in naïve CD5^hi^CD8^+^ T cells compared to the CD5^lo^ counterpart.(**D**) The dot plots of the significant KEGG gene set enrichment in the top 10 enriched pathways, reinforcing the presence of a poised phenotype for effector T cells in autoimmune diseases in naïve CD5^hi^CD8^+^ T cells. The data presented in (**A**) to (**D**) represent two experiments. In (**A**) to (**D**), n=10 mice per CD5^hi^CD8^+^ or CD5^lo^CD8^+^ group, with pooled spleen and PLN cells for sorting obtained from a total of 20 normoglycemia female NOD mice aged 6 to 8 weeks old. Significance is determined using a false discovery rate (FDR) adjusted *P*-value < 0.05 and validated using multiple testing correction method Benjamini-Hochberg correction.

To further characterize DEGs for their functional implications, we performed gene set analysis based on the gene ontology (GO). A gene-concept network by CD5^hi^ versus CD5^lo^ DEGs was associated with the three GO terms: external side of plasma membrane (GO:0009897), T cell activation (GO:0042110), and cytokine production involved in immune response (GO:0002367) (Supplementary Figure 2A). Among the notable findings, genes such as *Cd5*, *Cd83*, *Eomes*, *Xcl1*, *Tbx21*, *Il18rap*, *Klra1*, *Cxcr3*, and *Pdcd1* were significantly upregulated within CD5^hi^CD8^+^ T cells, with fold changes equal to or greater than 2. Specifically, within the GO term cytokine production involved in immune response, genes such as *Crlf2*, *Kit*, *F2rl1*, *Xcl1*, *Tbx21*, *Tnfrsf1b* and *Il18rap* showed upregulated expression patterns. Overall, these results provide insights into the potential mechanisms underlying T cell activation and cytokine production in CD5^hi^ population.

To thoroughly analyze the intrinsic kinetics and expression profiles of CD5^hi^CD8^+^ versus CD5^lo^CD8^+^ T cells under conditions resembling antigen encounter, we cross-referenced our findings with a dataset that delineates the functional relevance of ten gene clusters established in a prior study (47). This study identified core transcriptional signatures governing CD8^+^ T cell activation and effector/memory cell differentiation during infections, defining ten temporal gene-regulated patterns (clusters I∼X) in CD8^+^ T cells. Using a χ2 test to examine gene expression variations in these ten clusters between CD5^hi^ and CD5^lo^ cells (Table 1), we found that the CD5^hi^ subset has a significantly higher expression of genes linked to the initial cytokine or effector response (cluster I) and preparation for cell division (cluster II). Additionally, it exhibits a moderately higher expression of genes associated with cell cycle and division (cluster III), early effector and late memory (cluster V), short-term effector or memory (clusters IX), and late effector or memory (clusters X) in the CD8^+^ T cell response (Table 1 and Supplementary Figure 2B). Collectively, these data support that the CD5^hi^ T cells are better prepared for initial activation and late memory formation compared to the CD5^lo^ population.

**Table 1.**
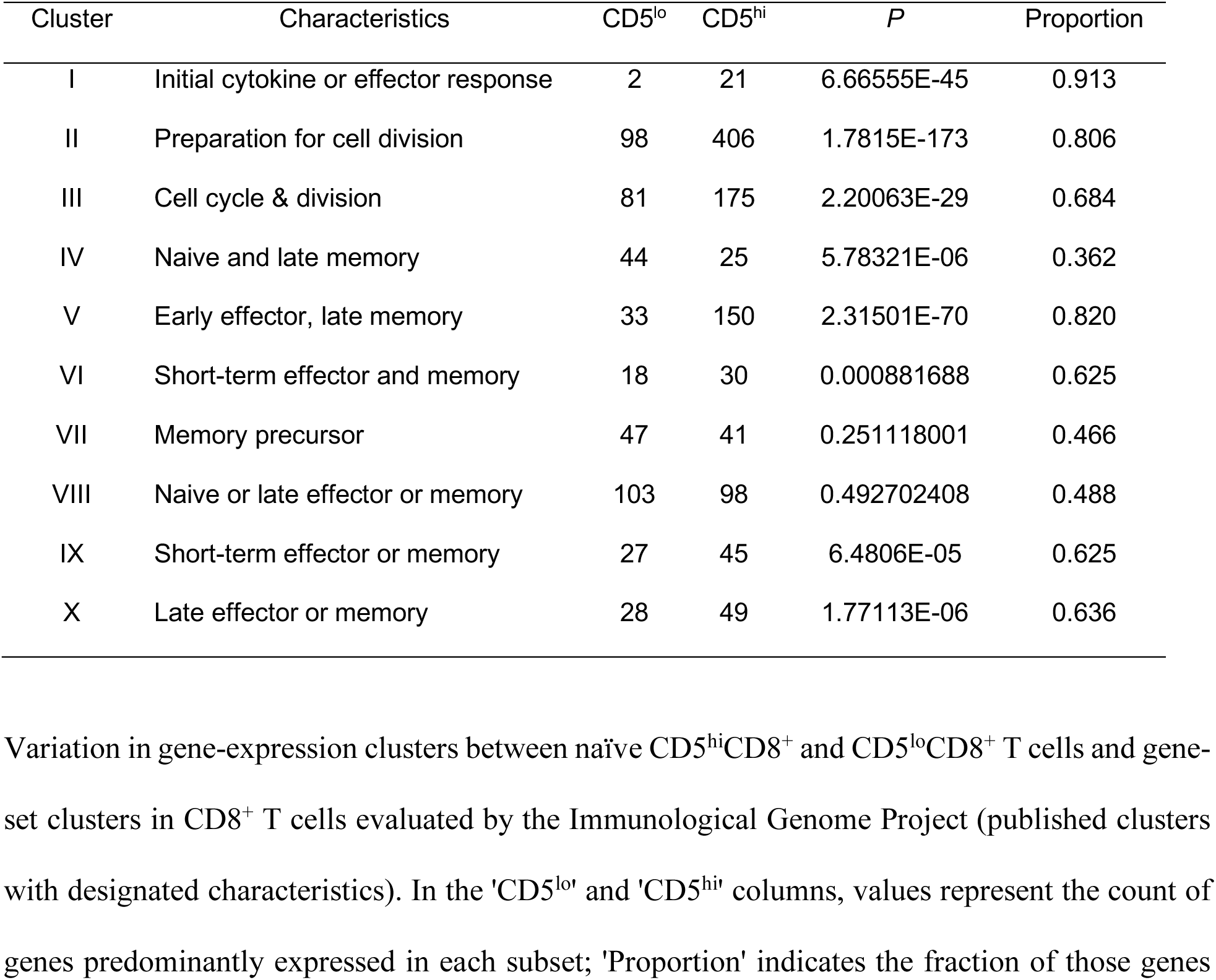
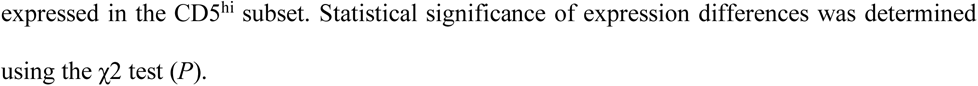
Naïve CD5^hi^CD8^+^ and CD5^lo^CD8^+^ T cells exhibit distinct expression in effector/memory T cell gene-expression clusters.

| Cluster | Characteristics | CD5<sup>lo</sup> | CD5<sup>hi</sup> | *P* | Proportion |
| --- | --- | --- | --- | --- | --- |
| I | Initial cytokine or effector response | 2 | 21 | 6.66555E-45 | 0.913 |
| II | Preparation for cell division | 98 | 406 | 1.7815E-173 | 0.806 |
| III | Cell cycle & division | 81 | 175 | 2.20063E-29 | 0.684 |
| IV | Naïve and late memory | 44 | 25 | 5.78321E-06 | 0.362 |
| V | Early effector, late memory | 33 | 150 | 2.31501E-70 | 0.820 |
| VI | Short-term effector and memory | 18 | 30 | 0.000881688 | 0.625 |
| VII | Memory precursor | 47 | 41 | 0.251118001 | 0.466 |
| VIII | Naïve or late effector or memory | 103 | 98 | 0.492702408 | 0.488 |
| IX | Short-term effector or memory | 27 | 45 | 6.4806E-05 | 0.625 |
| X | Late effector or memory | 28 | 49 | 1.77113E-06 | 0.636 |

In addition, we performed gene set enrichment analysis (GSEA) on DEGs using the Broad Institute’s GSEA desktop software. GSEA analysis indicate that CD5^hi^ cells were enriched for GO terms such as T cell activation (GO:0042110), external side of the plasma membrane (GO:0009897), positive regulation of immune system process (GO:0002684), regulation of catalytic activity (GO:0050790) and other GO terms with similar biological process (BP) functions (Figure 2C, Supplementary Figure 2C and Supplementary Table 3), similar to the results in Supplementary Figure 2A. Collectively, these results of cluster analysis (Table 1) and GSEA (Figure 2C) indicate that CD5^hi^CD8^+^ T cells in NOD mice possess a greater inherent potential for T cell activation and memory cell formation, which may contribute to the initiation and persistence of autoimmune diabetes.

Moreover, we analyzed DEGs by applying Kyoto Encyclopedia of Genes and Genomes (KEGG) database to further investigate whether CD5^hi^CD8^+^ subset is enriched in any potential disease-related pathways. Strikingly, two potential pathways stood out: the inflammatory bowel disease (IBD)- and type 1 diabetes mellitus-associated pathways (Figure 2D). The enrichment results for IBD and TID pathways suggest that naïve CD5^hi^CD8^+^ T cells with heightened self-reactivity have the intrinsic capacity to induce T1D and IBD. In summary, our data from gene-concept interactions, GSEA and KEGG pathways reveal distinct phenotypes in CD5^hi^CD8^+^ subset, indicating their potential for autoimmune predisposition even in their naïve state.

### CD5^hi^CD8^+^ and CD5^lo^CD8^+^ T cell subsets have distinct activation and proliferation levels

To examine whether variations in self-recognition, as indicated by CD5 expression, determine diabetogenic CD8^+^ T cell proliferation after TCR stimulation, we isolated naïve CD8^+^ T cells and subjected them to 2 days of dose-dependent TCR stimulation using various concentrations of anti-CD3 in combination with a consistent concentration of anti-CD28 (Figure 3A). Subsequently, we characterized phenotypic and functional distinctions within the 5% upper and lower CD5-expressing subsets of CD8^+^ T cells. Unlike the lower 5% CD8^+^ groups showing moderate change, the upper 5% CD8^+^ groups exhibited much more significant increase in CD5 levels in TCR stimulation dose-dependent manner (Figure 3B). The CD5^hi^CD8^+^ subset also showed higher p-CD3ζ abundance compared to CD5^lo^CD8^+^ subset across all TCR stimulation doses. Specifically, p-CD3ζ in CD5^hi^CD8^+^ peaked at 1.25 µg/ml anti-CD3 stimulation, then gradually declining with increased TCR stimulation doses, possibly due to its faster kinetic expression compared to other T cell activation signaling molecule p-Erk (Figure 3, C and D). Higher p-CD3ζ and p-Erk expressions in NOD CD5^hi^CD8^+^ cells compared to CD5^lo^CD8^+^ counterparts, consistent across TCR stimulation doses, align with the previous findings in B6 mice (2, 5). The upper 5% CD5^hi^ cells consistently exhibited elevated levels of T cell activation markers CD69 and CD44 (Figure 3, E and F) and showed dose-dependent enhanced proliferation under TCR stimulation (Figure 3G), along with their increased CD127 level (Figure 3H), suggesting heightened capabilities in memory T cell homeostasis.

**Figure 3.**
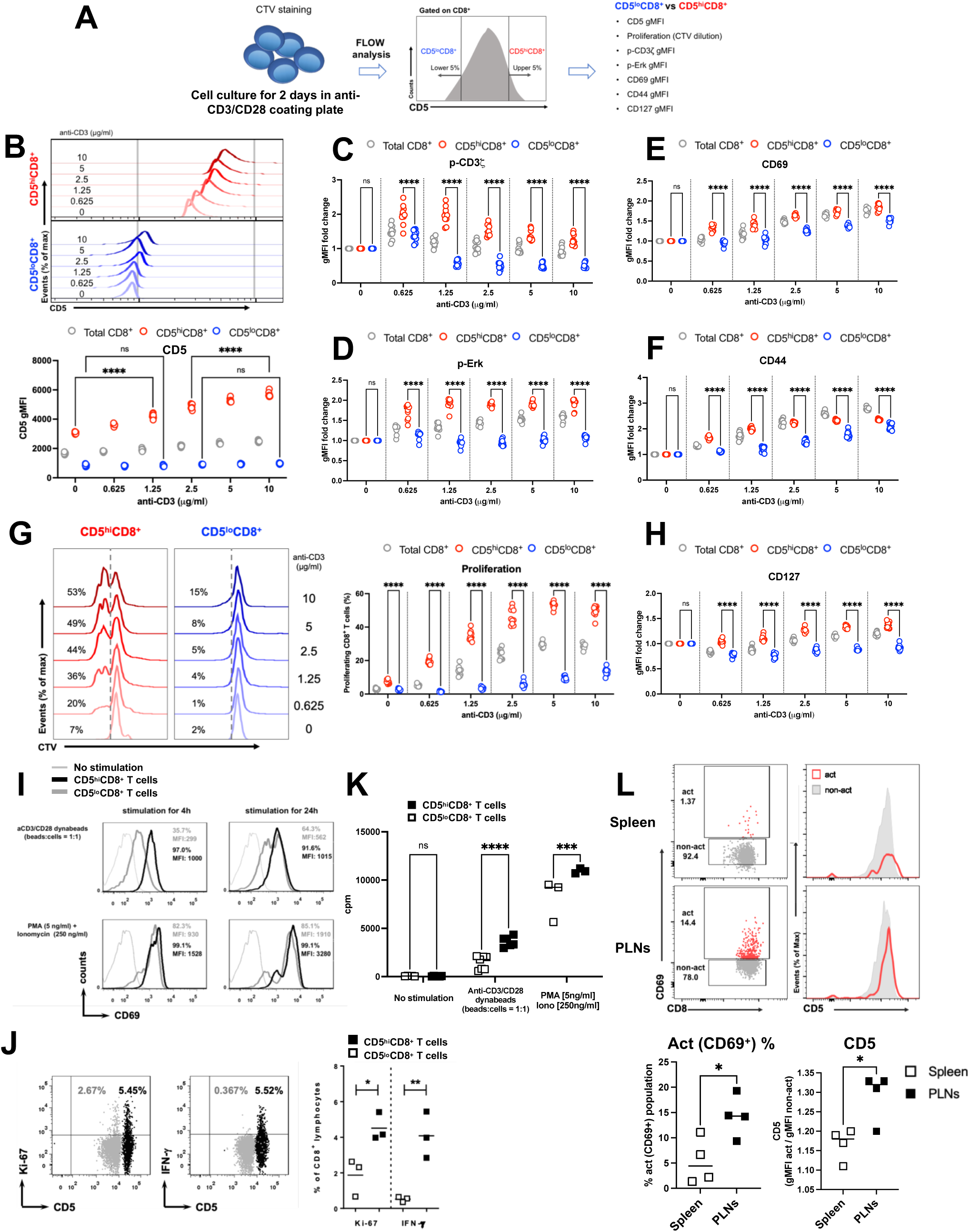
CD5 expression distinguishes CD8^+^ T cell subsets with differential activation and proliferation. (**A**-**K**) Naïve CD8⁺ T cells were isolated from the spleen of 6–8-week-old normoglycemic female NOD mice and stimulated with the indicated conditions. (**A**) Diagram depicting experimental setup using varying concentrations of anti-CD3 (0–10 μg/ml) and anti-CD28 (2 μg/ml; 0 μg/ml for control) for 2 days. (**B**, **G** upper/left panels) Representative flow cytometry plots, with statistical summaries shown in (**B**-**H**). Comparison of CD5 (**B**), fold change of p-CD3ζ (**C**), p-Erk (**D**), CD69 (**E**), CD44 (**F**), proliferation levels (CTV dilution %) (**G**) and fold change of CD127 expression (**H**) between the upper and lower 5% of CD5-expressing CD8⁺ T cells (CD5^hi^CD8⁺ and CD5^lo^CD8⁺, respectively). Fold change values in (**C**-**F**, **H**) were normalized to each group’s baseline at 0 μg/ml anti-CD3 stimulation, where total CD8⁺ T cells, CD5^hi^CD8⁺ and CD5^lo^CD8⁺ were each set to 1. (**I**) Histogram comparing CD69 expression between the upper and lower 10% of CD5-expressing CD8⁺ T cells (CD5^hi^CD8⁺ and CD5^lo^CD8⁺) after stimulation with anti-CD3/CD28 dynabeads or PMA plus Ionomycin for specified durations. (**J**) Comparison of Ki-67 expression and IFN-γ production between CD5^hi^CD8⁺ and CD5^lo^CD8⁺ T cells.,(**K**) Proliferation of CD5^hi^CD8⁺ and CD5^lo^CD8⁺ T cells assessed by [methyl-3H] thymidine incorporation (cpm; counts per minute) after stimulation with the indicated amounts of anti-CD3/CD28 dynabeads or PMA plus Ionomycin for 2 days. (**L**) Comparison of CD5 gMFI in CD8^+^CD69^+^ (upper gate of flow dot plot) to that in CD8^+^CD69^-^ (lower gate of flow dot plot) population by histogram (CD8^+^CD69^+^ in solid red line; CD8^+^CD69^-^ in grey), across spleen and PLNs from normoglycemic female NOD mice aged 6 to 8 weeks. The statistical results of the percentages of activated (CD8^+^CD69^+^) population (left) and the ratio of activated to non-activated CD5 gMFI (right) are shown. Data represent mean ± SD. The data presented in (**B**) to (**H**) represent two experiments. The sample size was n = 10 (**B**-**H**), n = 3 (**J**), n = 3-7 (**K**), and n = 4 (**L**) per group. \**P* < 0.05, \*\**P* < 0.01, \*\*\**P* < 0.001, \*\*\*\**P* < 0.0001, by two-way ANOVA (**B**)-(**K**) and unpaired, two-tailed t test (**L**).

In Figure 3, C-H, we exclusively evaluated the phenotypic and functional differences between CD5^hi^ and CD5^lo^ subsets from activated bulk CD8^+^ T cells following TCR stimulation. To investigate whether these distinctions post TCR activation stem from intrinsic differences between CD5^hi^ and CD5^lo^ cells, we specifically sorted the upper and lower 10% of CD5-expressing CD8^+^ T cells from prediabetic NOD mice aged 6 to 8 weeks, aiming to ascertain whether sorted naive CD5^hi^ and CD5^lo^ T cells maintain these intrinsic differences in their immune responses after TCR or downstream signaling stimulation (Figure 3, I-K). The results revealed that CD5^hi^ cells, after TCR stimulation, displayed increased levels of the T cell activation marker CD69 (Figure 3I), accompanied by higher Ki-67 expression and IFN-γ production (Figure 3J), reflecting their intrinsic distinctions in heightened effector and proliferation functions. Notably, in our T cell stimulation experiment comparing CD5^hi^ and CD5^lo^CD8^+^ T cells treated with anti-CD3/CD28 dynabeads, we observed superior activation potential in CD5^hi^ cells (Figure 3I). After 4 hours of stimulation, CD69 expression in CD5^hi^ cells was significantly higher than in CD5^lo^ cells, with a sharp distinction. As the stimulation time extended to 24 hours, CD69 levels in CD5^hi^ compared to CD5^lo^ remained higher, although the difference reduced, and CD69 expression in both populations became closer. In comparison, with PMA/Ionomycin treatment bypassing TCR signaling initiation, there were almost no distinctions in CD69 levels between CD5^hi^ and CD5^lo^ populations, implying cell activation differences between CD5^hi^ and CD5^lo^CD8^+^ T cells primarily through TCR-proximal signaling. The DNA analog [methyl-3H] thymidine incorporation assay further confirmed the heightened proliferation capacity in CD5^hi^ subset compared to CD5^lo^ subset (Figure 3K). Additionally, we compared CD5 levels between CD8^+^CD69^+^ and CD8^+^CD69^-^ populations in both the spleen and PLNs. In PLNs, a lymphoid tissue enriched with autoantigens, CD69^+^CD8^+^ T cells were skewed toward expressing higher CD5 levels compared to the counterparts in spleen (Figure 3L), implying that cells with high CD5 level display elevated activation potential upon encountering antigen. In summary, these results indicate that diabetogenic CD8^+^ T cells exhibiting increased activation and proliferation profiles in NOD mice tend to have elevated CD5-associated auto-immune responses.

### The CD5 expression level in naïve CD8^+^ T cells is positively correlated with diabetes susceptibility in NOD mice

We further explored potential differences in *in vivo* immune response of CD5^hi^ versus CD5^lo^CD8^+^ T cells, especially their abilities to induce insulitis and diabetes through an adoptive T cell transfer experiment in NOD model. Following the similar sorting method as in Figure 3, I-K, the naïve CD5^hi^CD8^+^ and CD5^lo^CD8^+^ cells from prediabetic NOD mice were then administered separately to immunodeficient NOD Rag1^-/-^ recipients via intraperitoneal injection. The diabetes incidence in Rag1^-/-^ mice was weekly monitored after transfer. Strikingly, the CD5^hi^-transferred group exhibited markedly greater diabetes onset kinetics compared to CD5^lo^ group, with the appearance of diabetes as early as 4 weeks post-transfer. All 10 mice in CD5^hi^ group experienced earlier diabetes onset, in contrast to the CD5^lo^-transferred group, where only 4 out of 11 mice exhibited diabetes after approximately 20 weeks post-transfer (Figure 4A, left panel). Moreover, histological analysis and insulitis scoring of pancreata after the 4-week transfer period revealed heightened severity of insulitis, characterized by profound lymphocytic infiltration in the CD5^hi^CD8^+^ compared to CD5^lo^CD8^+^ group (Figure 4A, right panel). This suggests that CD5^hi^CD8^+^ cells possess more potent diabetogenic properties during the immune responses in autoimmune diabetes.

**Figure 4.**
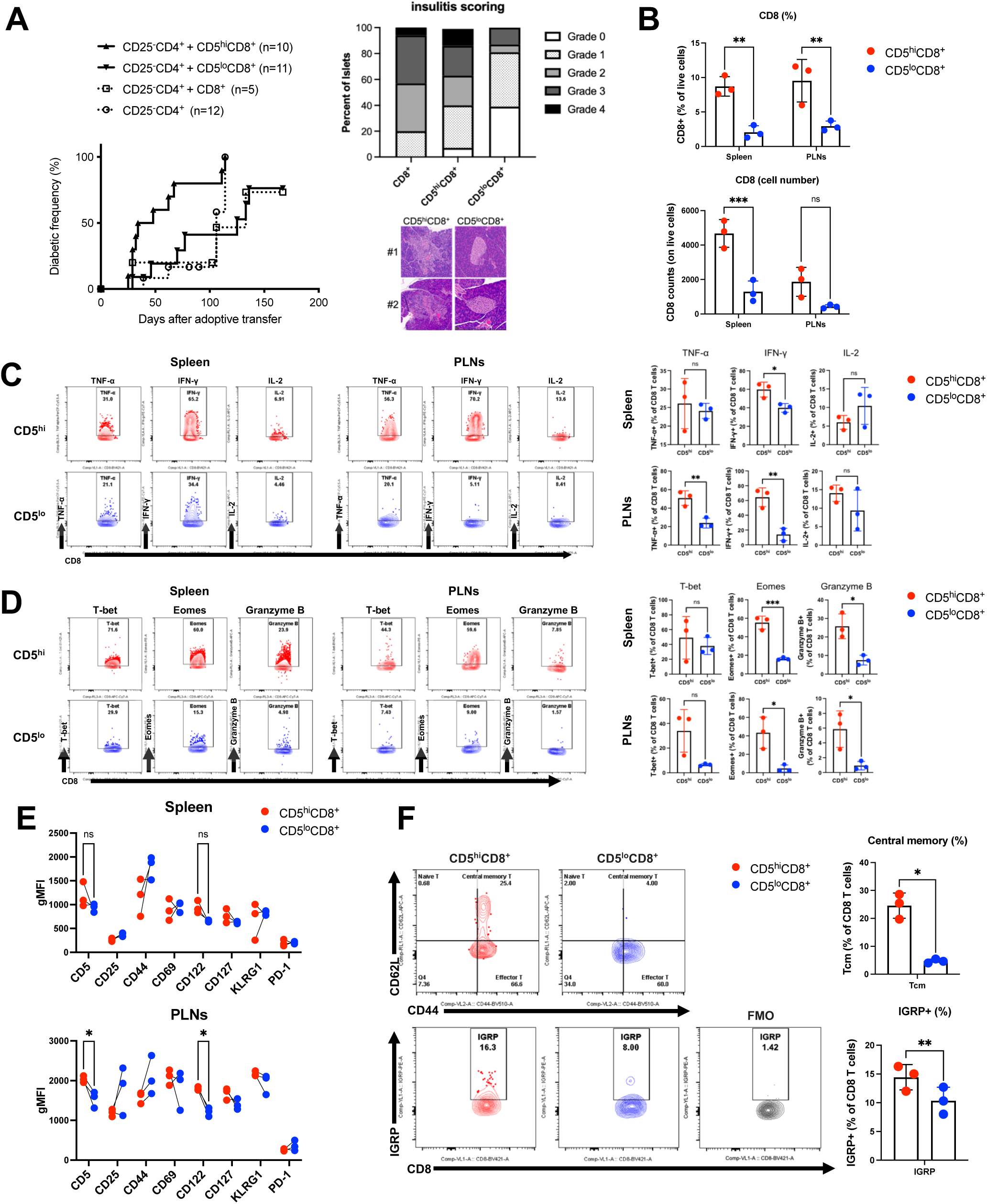
CD5^hi^CD8^+^ T cells-transferred group exhibits severe insulitis and disease phenotype. (**A**) Diabetes incidence (left) in NOD Rag1^-/-^ mice transferred with naïve CD5^hi^CD8^+^, CD5^lo^CD8^+^, or CD8^+^ T cells, combined with CD4^+^ T cells, respectively. The islet histological analyses and insulitis scoring (right) for the mice after a 4-week transfer in the respective transferred groups showed a higher diabetes frequency in the CD5^hi^CD8^+^ group compared to the CD5^lo^CD8^+^ group. The cells were sorted from the 10% upper and lower of CD5 expression of naïve CD8^+^ T cells (CD5^hi^CD8^+^ and CD5^lo^CD8^+^ T cells) from the spleen and PLNs 6-8-week-old normoglycemic female NOD mice and adoptively transferred to 6-8- week-old female NOD Rag1^-/-^ recipients via i.p. injection. Urine glucose concentrations of the four groups of mice were monitored weekly for diabetes incidence. Diabetes was defined as glycosuria > 500 mg/dl in two consecutive tests. Insulitis scoring criteria can be found in the supplementary materials and methods. (**B**) Percentages and cell numbers of CD8^+^ T cells in spleen and PLNs, analyzed from CD5^hi^CD8^+^ and CD5^lo^CD8^+^ cells-transferred group, respectively, after 4-week adoptive transfer. (**C**) Percentages of TNF-α, IFN-γ and IL-2 production in CD5^hi^CD8^+^ and CD5^lo^CD8^+^ group, respectively, from spleen or PLNs, analyzed by flow cytometry, with representative flow dot plots (left) and statistical results (right). (**D**) Percentages of T-bet, Eomes and Granzyme B-expressing in CD5^hi^CD8^+^ and CD5^lo^CD8^+^ group, respectively, from spleen or PLNs, analyzed by flow cytometry, with representative flow dot plots (left) and statistical results (right). (**E**) Comparison of differential gMFI expression of CD5, CD25, CD44, CD69, CD122, CD127, KLRG1 and PD-1 between CD5^hi^CD8^+^ and CD5^lo^CD8^+^ group from spleen or PLNs. (**F**) Comparison of percentages of central memory T cells (defined by CD44^hi^CD62L^hi^) (upper) and positive IGRP-tetramer staining (lower) between CD5^hi^CD8^+^ and CD5^lo^CD8^+^ groups from PLNs, with representative flow dot plots (left) and statistical results (right). Representative flow cytometry plots include FMO negative control for accurate IGRP-tetramer gating. Data represent mean ± SD. Sample size was n = 5-11 (**A**), and n = 3 (**B**-**F**) per group. \**P* < 0.05, \*\**P* < 0.01, \*\*\**P* < 0.001, by log-rank test (**A**), by two-way ANOVA (**B**) and (**E**) and unpaired, two-tailed t test (**C**), (**D**) and (**F**).

Moreover, flow cytometry analysis revealed a significant increase in both the percentage and the number of CD8 T cells in the spleen and PLNs of the CD5^hi^-transferred group compared to their CD5^lo^ counterparts (Figure 4B), implying increased expansion and activation/effector potential by the CD5^hi^ subset, consistent with the observations in *in vitro* proliferation assay (Figure 3K). Additionally, the CD5^hi^ group exhibited higher levels of IFN-γ and a trend toward increased TNF-α, while IL-2 production did not show a significant difference (Figure 4C) and higher percentages of CD8^+^ T cells expressing the transcription factors T-bet, Eomes and the cytotoxic molecule Granzyme B (Figure 4D). In characterizing other key immune markers between CD5^hi^ and CD5^lo^ groups, as shown in Figure 1, E and F, we observed that CD5^hi^ cells- transferred group in PLNs exhibited high level of CD122 but not in spleen, while also maintaining high CD5 expression (Figure 4E). Furthermore, we assessed the composition of central memory T cells, defined by the CD44^hi^CD62L^hi^ phenotype, and quantified the percentages of CD8^+^ T cells that recognize islet-specific antigen IGRP_206–214_ by tetramer staining (IGRP^+^CD8^+^ T cells) in the PLNs (Figure 4F). This analysis underscores that central memory T cell population and the frequency of islet autoantigen-specific CD8 T cells are higher in the CD5^hi^ transferred subset within the PLNs, implying more robust immune responses initiated by the CD5^hi^ cells.

### The intrinsic high CD5-linked self-reactivity of CD8^+^ T cells attributed to thymic positive selection contributes to the peripheral poised autoimmune phenotypes

Based on our results, phenotypic and functional heterogeneity was observed within naive CD8^+^ T cells with different CD5 level (Figure 1 and 2), and the further sorted CD5^hi^ and CD5^lo^ cells displayed differential effector/memory functions (Figure 3 and 4). To validate that the differential autoimmune T cell responses in CD5^hi^ and CD5^lo^CD8^+^ T cells are linked to heterogeneity in TCR self-reactivity during positive selection, we examined whether the unique TCR basal signal strength and IL-2 production differs in the lower and upper CD5-expressing thymocytes in NOD mice. Indeed, from the immature CD4^−^CD8^−^ double-negative (DN), CD4^+^CD8^+^ double-positive (DP), to CD4^−^CD8^+^ (CD8SP) stages in 6-8-week-old NOD thymocytes, the high self-reactivity CD5^hi^ thymocytes exhibited elevated p-CD3ζ, p-Erk and IL-2 expression, supporting their characteristics of survival and proliferation (Figure 5A and Supplementary Figure 3, A and B). These findings are consistent with previous report (2) and may explain a higher CD8^+^ T cell expansion observed in CD5^hi^CD8^+^ cell-transferred recipients compared to CD5^lo^CD8^+^ cell-transferred recipients in the adoptive transfer experiment (Figure 4B). We further explored the potential influence of thymic selection on peripheral T cell responses by using the islet antigen-specific 8.3 TCR transgenic NOD8.3 mice, characterized by higher CD5 expression, compared to that of NOD mice, and thus an enhanced level of self-recognition. NOD8.3 compared to NOD mice consistently showed elevated CD5 expression from DP to CD8SP stages (Figure 5B), and a significant increase in cell number in the CD8SP subset (Supplementary Figure 3, A and C), suggesting an elevated positive selection in NOD8.3 thymocytes. NOD8.3 mice, with elevated CD5 expression linked to basal TCR signal strength (10), showed significantly higher p-CD3ζ level in DP cells compared to NOD mice counterparts (Figure 5C and Supplementary Figure 3D). The increase of p-CD3ζ level in DP cells positively correlates with the alterations in thymocyte development reflected in increased cell number and percentage in the CD8SP subset transitioning from the DP stage of NOD8.3 mice (Figure 5D and Supplementary Figure 3C). Additionally, DN thymocytes, marked by CD44 and CD25, undergo crucial β-selection in DN3 (CD44^-^CD25^+^) and DN4 (CD44^-^CD25^-^), emphasizing the importance of DN stage in thymocyte development (48–51). NOD8.3 mice showed significantly higher cell numbers in DN4 (Figure 5E and Supplementary Figure 3E), suggesting a selection advantage in these stages due to bypassing the need for TCRβ rearrangement occurred in NOD cells.

**Figure 5.**
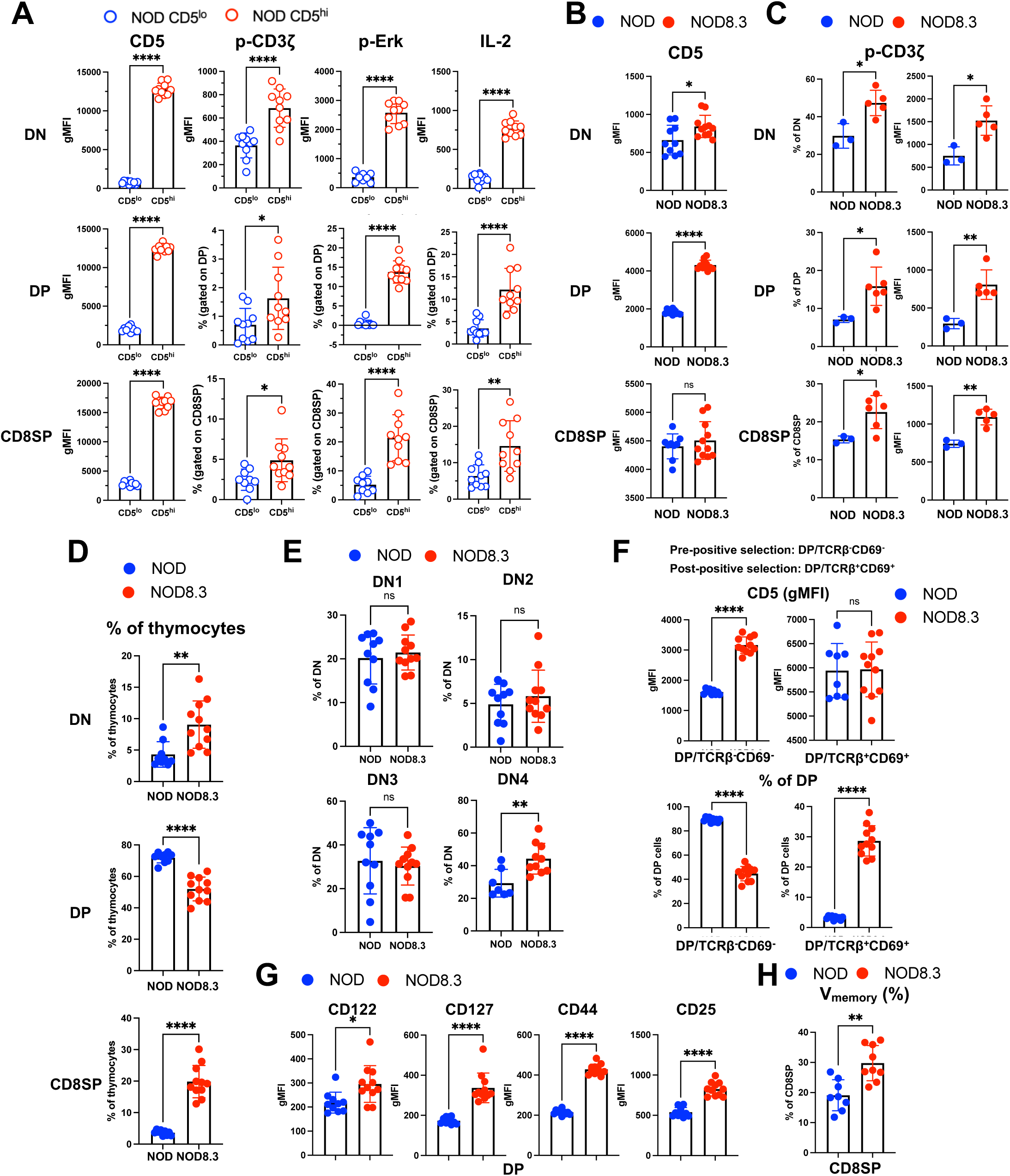
The high CD5-linked TCR basal signals positively correlate with thymic selection in autoimmune-poised NOD8.3 mice. (**A**) CD5, p-CD3ζ, p-Erk and IL-2 expression levels compared between the lower 5% (hollow blue) and upper 5% (hollow red) CD5 expression in thymocytes of normoglycemia female NOD mice aged 6 to 8 weeks old across DN, DP and CD8SP stages. (**B**-**H**) Comparison of various marker expression levels between thymocytes of normoglycemia female NOD (solid blue) and NOD8.3 (solid red) mice aged 6 to 8 weeks old across different selection stages. (**B**) CD5 gMFI in NOD and NOD8.3 thymocytes across DN, DP and CD8SP stages. (**C**) Percentages of p-CD3ζ (left) and the corresponding gMFI of p-CD3 (right) compared between NOD and NOD8.3 thymocytes in the indicated stages DN, DP and CD8SP. (**D**) Percentages of DN, DP and CD8SP composition compared between NOD and NOD8.3 thymocytes. (**E**) Percentages of indicated DN stages (DN1, DN2, DN3 and DN4) in the DN subset compared between NOD and NOD8.3 mice. DN stages are defined as follows: DN1 (CD44^+^CD25^-^), DN2 (CD44^+^CD25^+^), DN3 (CD44^-^CD25^+^) and DN4 (CD44^-^CD25^-^), as depicted in the gating shown in Supplementary Figure 3A, middle panels. (**F**) CD5 gMFI (upper) and corresponding percentages (lower) in the pre-positive selection (TCRβ^-^CD69^-^) and post-positive selection (TCRβ^+^CD69^+^) populations in DP thymocytes compared between NOD and NOD8.3 mice. The presence of higher TCR basal signals (indicated by higher CD5 gMFI) in NOD8.3 in the pre-positive selection stage leads to an increased percentage of TCRβ^+^CD69^+^ cells in the post-selection stage compared to NOD mice, as shown in Supplementary Figure 3F. (**G**) gMFI of CD122, CD127, CD44 and CD25 assessed in DP thymocytes compared between NOD and NOD8.3 mice. (**H**) Percentages of virtual memory (V_memory_) cells in CD8SP thymocytes compared between NOD and NOD8.3 mice. Data represent mean ± SD. The data presented in (**B**) to (**H**) represent two experiments. The sample size was n = 10 (**A**), n = 10-11 (**B**), n = 3-6 (**C**), n = 9-11 (**D**), n = 7-11 (**E**), n = 8-11 (**F**) n = 10-11 (**G**) and n = 8-9 (**H**) per group. \**P* < 0.05, \*\**P* < 0.01, \*\*\*\**P* < 0.0001, by unpaired, two-tailed t test (**A**- **H**).

Positive selection in the DP stage was further characterized by analyzing CD69 and TCRβ levels. In the pre-positive selection population (DP/TCRβ^-^CD69^-^), NOD8.3 cells showed significantly higher CD5 expression compared to NOD cells (Figure 5F, upper panel, and Supplementary Figure 3F), implying an enhanced TCR signal perception at the initiation of DP positive selection (52). Consistently, the frequency of CD69^+^TCRβ^+^ cells, representing post-positive-selection thymocytes (53), significantly increased in NOD8.3 compared to NOD mice (Figure 5F, lower panel), predisposing to an accelerated accumulation of CD8SP cells (Figure 5D and Supplementary Figure 3C). Additionally, to investigate whether the increased accumulation of CD8SP cell number in NOD8.3 mice is associated with elevated expressions of cytokine receptors crucial for T cell survival in DP stage, we assessed the expression levels of CD122, CD127, CD44 and CD25 in DP thymocytes. Indeed, these receptors showed higher levels in NOD8.3 compared to NOD DP thymocytes (Figure 5G), suggesting their contribution to cell survival into the CD8SP stage. Remarkably, NOD8.3 mice showed an increased percentage of memory-like CD122^hi^CD44^hi^ cells (V_M_; virtually memory) in CD8SP compared to NOD mice (Figure 5H and Supplementary Figure 3G), implying a direct transition to memory T cells upon entering peripheral tissues (54). Furthermore, CD5 expression patterns were compared across thymocyte stages among different NOD strains including NOD, BDC2.5 and NOD8.3 mice. NOD8.3 thymocytes consistently showed higher CD5 levels compared to two other strains from the DN to DP stages, and notably, in the CD8SP stage, NOD8.3 cells exhibited the highest CD5 expression, followed by NOD and then BDC2.5 cells (Supplementary Figure 4A). Interestingly, NOD8.3 cells displayed enhanced proliferation (Supplementary Figure 4, B and C), suggesting a potential link between perceived TCR basal signals during thymocyte selection and peripheral lymphocyte proliferation.

### Downregulation of TCR signaling strength by transgenic phosphatase Pep expression does not confer the protection in dLPC/NOD8.3 mice

Based on the insights gained from thymocyte development in NOD, BDC2.5 and NOD8.3 mice, where the TCR signal strength influences intrinsic T cell response and subsequent effector function, we investigated whether the manipulation on TCR basal signal strength influences autoimmune potential of peripheral T cells in NOD mice. To address this issue, we utilized two transgenic mouse model, dLPC/NOD and dLPE/NOD mice, which express a T cell-specific phosphatase Pep to downregulate proximal TCR signaling (27). A significant decrease in autoimmune diabetes, in terms of a delayed onset, less insulitis severity and lower incidence of disease, was found in dLPC/NOD and dLPE/NOD mice compared to NOD controls (27). As expected, transgenic Pep expression in dLPC/NOD mice delayed the onset of diabetes and reduced insulitis. However, Pep overexpression in dLPC/NOD8.3 mice did not provide protection (Figure 6A), suggesting that intrinsic high CD5-associated self-reactivity in NOD8.3 T cells overrides the transgenic Pep-mediated protection observed in dLPC/NOD mice. Moreover, overexpression of Pep in dLPC/NOD and dLPC/NOD8.3 mice did not change the CD5 expression of T cells in the spleen compared to their controls, NOD and NOD8.3, respectively. Interestingly, we observed a transgenic Pep-mediated CD5 downregulation in CD8^+^ T cells in PLNs, enriched with islet-specific autoantigens, of dLPC/NOD mice compared to NOD controls (Figure 6B), suggesting that the transgenic Pep-mediated TCR signaling strength modulates CD5 expression in a context-specific manner. However, CD5 expression level in CD8^+^ T cells in PLNs of dLPC/NOD8.3 was similar to NOD8.3 mice (Figure 6B), suggesting that NOD8.3 T cells with an intrinsic high CD5- associated self-reactivity are more resistant to transgenic Pep-mediated change in TCR signaling and CD5 expression. These observations are further supported by our data that the CD5-correlated basal TCR signaling molecule, p-CD3ζ, was attenuated in dLPC/NOD mice but not in dLPC/NOD8 .3 mice, both in the spleen and PLNs (Figure 6C). Collectively, our results suggest that CD8^+^ T cells with intrinsic high CD5-linked self-reactivity are more resistant to transgenic Pep-mediated TCR signaling downregulation and sustain the diabetogenic susceptibility in NOD8.3 mice.

**Figure 6.**
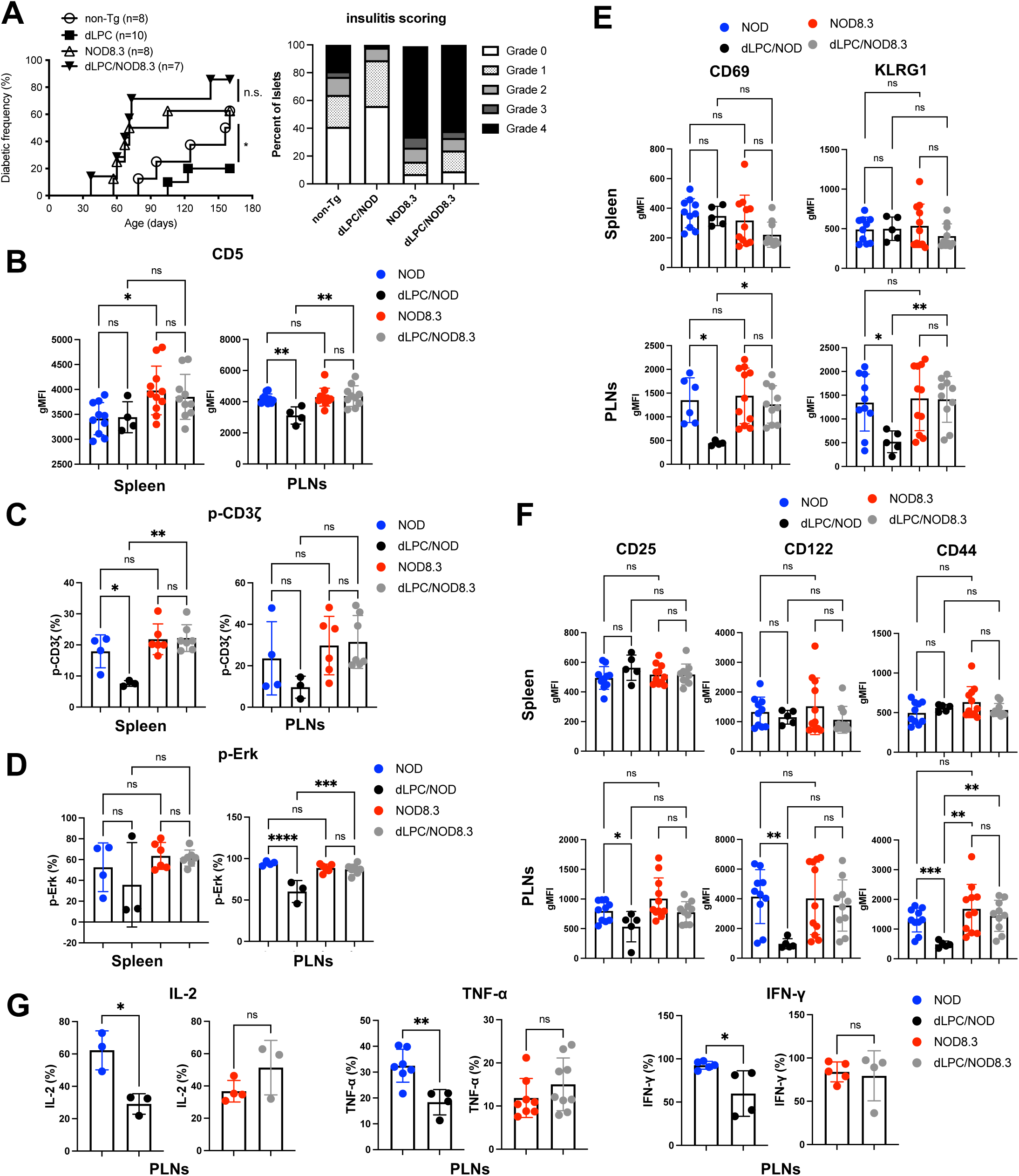
The protective effect of transgenic Pep on diabetogenesis is negated in dLPC/NOD8.3 mice. (**A**) Spontaneous diabetes incidence (left) and insulitis scoring (right) of female non-Tg NOD, dLPC/NOD, NOD8.3 and dLPC/NOD8.3 mice. Insulitis scoring was performed on the islets from the pancreata of 6-8-week-old mice before the onset of diabetes in each group. Insulitis scoring criteria can be found in the supplementary materials and methods. Urine glucose concentrations of mice were monitored weekly for spontaneous diabetes incidence. Diabetes was defined as glycosuria > 500 mg/dl at two consecutive tests. (**B**-**G**) Characterization of various aspects of TCR signal strength and effector/memory T cell phenotypes among female normoglycemia non-Tg NOD (blue), dLPC/NOD (black), NOD8.3 (red) and dLPC/NOD8.3 (grey) mice at the age of 6 to 8 weeks old. (**B**) CD5 gMFI of CD8^+^ T cells in the spleen (left) and PLNs (right). (**C**) Percentage of p-CD3ζ^+^CD8^+^ T cells in the spleen (left) and PLNs (right). (**D**) Percentage of p-Erk^+^CD8^+^ T cells in the spleen (left) and PLNs (right). (**E**) gMFI of effector T cell markers CD69 and KLRG1 in CD8^+^ T cells from the spleen (upper) and PLNs (lower). (**F**) gMFI of memory T cell markers CD25, CD122 and CD44 in CD8^+^ T cells from the spleen (upper) and PLNs (lower). (**G**) Percentage of IL-2^+^-, TNF-α^+^-, and IFN-γ^+^CD8^+^ T cells in PLNs. Data in (**B**) to (**G**) represent mean ± SD. The data presented in (**B**) to (**G**) represent two experiments. The sample size was n = 7-10 (**A**), n = 4-11 (**B**), n = 3-7 (**C**), n = 3-7 (**D**), n = 5-11 (**E**), n = 5-11 (**F**) and n = 3-9 (**G**) per group. \**P* < 0.05, \*\**P* < 0.01, \*\*\**P* < 0.001, \*\*\*\**P* < 0.0001, by log-rank test (**A**), by one-way ANOVA (**B**-**F**) and unpaired, two-tailed t test (**G**).

Consistently, isolated naïve CD8^+^ T cells (Supplementary Figure 5 A, left panel) from high CD5-expressing NOD8.3 mice exhibited superior proliferation compared to cells from NOD mice upon anti-CD3/CD28 or PMA/Iono stimulation (Supplementary Figure 5A, middle and right panel, respectively). Moreover, transgenic Pep expression attenuated CD8^+^ T cell proliferation in both dLPC/NOD and dLPE/NOD mice compared to NOD controls (Supplementary Figure 5B). However, transgenic Pep expression did not alter cell proliferation in dLPE/NOD8.3 T cells when stimulated with the autoantigen-specific self-peptide IGRP_206-214_ (Supplementary Figure 5C), suggesting that a high TCR basal signal conferred by elevated intrinsic self-reactivity overrides the transgenic Pep-mediated reduction of T cell proliferation. Further analysis of p-Erk level also showed a significant decrease in dLPC/NOD compared to NOD mice. Similarly, transgenic Pep did not attenuate the p-Erk level in dLPC/NOD8.3 mice (Figure 6D), consistent with the observation of p-CD3ζ level in these Pep transgenic mice (Figure 6C). Additionally, CD69 and KLRG1 (Figure 6E) and other activation markers including CD44 and CXCR3 (Supplementary Figure 5D) were decreased in dLPC/NOD or dLPE/NOD mice compared to NOD controls. Similar to previous observations, transgenic Pep-mediated attenuation of T cell activation was abolished in dLPC/NOD8.3 mice, supporting again that an intrinsic high CD5-linked self-reactivity is able to confer the impact by transgenic Pep-mediated TCR signaling change.

In addition, transgenic Pep expression downregulated CD25 level in T cells of PLNs, not in spleen, in dLPC/NOD mice compared to NOD mice, whereas there was no significant change in CD25 expression between dLPC/NOD8.3 and NOD8.3 mice (Figure 6F). Moreover, memory T cell homeostasis molecules, such as CD122 and CD44, were also reduced in T cells of PLNs, not in spleen, in dLPC/NOD mice compared to NOD mice (Figure 6F and Supplementary Figure 5D), but were indistinguishable between dLPC/NOD8.3 and NOD8.3 mice (Figure 6F). Notably, transgenic Pep expression in dLPC/NOD mice significantly reduced production of effector T cell cytokines, such as IL-2, TNF-α and IFN-γ in CD8^+^ T cells compared to NOD mice (Figure 6G and Supplementary Figure 5E). Nevertheless, transgenic Pep-mediated reduction in cytokine production was abolished in dLPC/NOD8.3 mice (Figure 6G). Collectively, our results indicate that CD5-associated high self-reactivity is able to abolish transgenic Pep-mediated attenuation of effector/memory T cell functions in NOD8.3 mice.

### Analysis of the CD5^hi^ population identifies TCR repertoires associated with autoimmune diseases

To investigate whether pathogenic feature in CD5^hi^CD8^+^ population reflects in their TCR repertoire, we divided naïve CD8^+^ T cells into CD5^hi^CD8^+^ and CD5^lo^CD8^+^ groups, conducted bulk RNA sequencing on two groups, and applied the MiXCR pipeline to analyze TCR repertoire composition in CD5^hi^ and CD5^lo^ groups (Figure 7A and Supplementary Figure 6). Analysis of T-cell receptor-α chain (*TRAV* and *TRAJ*) and T-cell receptor-β chain (*TRBV* and *TRBJ*) gene usages in CD5^hi^CD8^+^ and CD5^lo^CD8^+^ T cells revealed increased utilization of *TRAV13-2*, *TRAV16D- DV11*, *TRAV3-3*, *TRAJ32*, *TRBV14*, *TRBV16*, *TRBJ2-1* and *TRBJ2-7* in CD5^hi^CD8^+^ cells, with the exception of *TRAV9-1*, which exhibited lower utilization in CD5^hi^CD8^+^ T cells (Figure 7, B-E). The CDR3 loops of TRA (CDR3α) and TRB (CDR3β), crucial for antigen recognition by TCRs on T cells, are shaped by the diverse recombination events that assemble TRA VJ gene segments and TRB V(D)J gene segments (55). The distinct utilization of V or J gene families of TCRα and TCRβ, as shown in Figure 7, B-E, may lead to variable CDR3 characteristics between CD5^hi^CD8^+^ and CD5^lo^CD8^+^ T cells. Additionally, emerging evidence suggests a potential association of shorter CDR3 length in T1D patients (56). Therefore, we analyzed whether the CD5^hi^CD8^+^ cells have similar pathogenic CDR3 patterns. Interestingly, examination of CDR3α and CDR3β length distribution in the TCR repertoire revealed a bias toward shorter length in the CD5^hi^ compared to the CD5^lo^ group, particularly with a significantly shorter CDR3β length (Figure 7F). Shared TCR pools were reported to be enriched in clonotypes with fewer insertions in CDR3 (57). To confirm this, the overlapped clonal count matrix for CDR3α or CDR3β motif was calculated among all groups (Figure 7G), indicating increased similarity between the CD5^hi^ groups based on its own pairing comparison. Strikingly, the IGRP-recognizing CDR3 motif was found in CDR3α of the CD5^hi^ group (Figure 7H), which has been identified in the TCR repertoire of T1D patients (58). Moreover, analysis of CDR3α and CDR3β amino acid characteristics revealed a hydrophobicity enrichment in TRA of CD5^hi^CD8^+^ cells (Figure 7I), consistent with a previous study demonstrating that the generation of high self-reactive T cells was promoted by hydrophobic residues within the CDR3 region (59). Furthermore, in alignment with a previous study illustrating greater diversity in TCR repertoires from T1D patients than in healthy donors (56), CD5^hi^CD8^+^ cells exhibited higher diversity in clonotypes compared to their CD5^lo^ counterpart (Figure 7J). Additionally, our results align with a previous study that employed high-throughput TCR-seq analyses in mice, identifying highly shared CDR3 sequences among mice characterized by restricted V and J segment usage (60). These CDR3 sequences, exhibiting lower nucleotide insertions and shorter lengths, were more abundant, particularly in individuals displaying autoimmune and allograft- related reactions, further affirming their association with self-antigen reactivity. In conclusion, these results suggest that CD5^hi^CD8^+^ T cells, with elevated self-reactivity and increased clonal diversity, are intrinsically linked to the utilization of specific V and J genes within CD5^hi^ cells.

**Figure 7.**
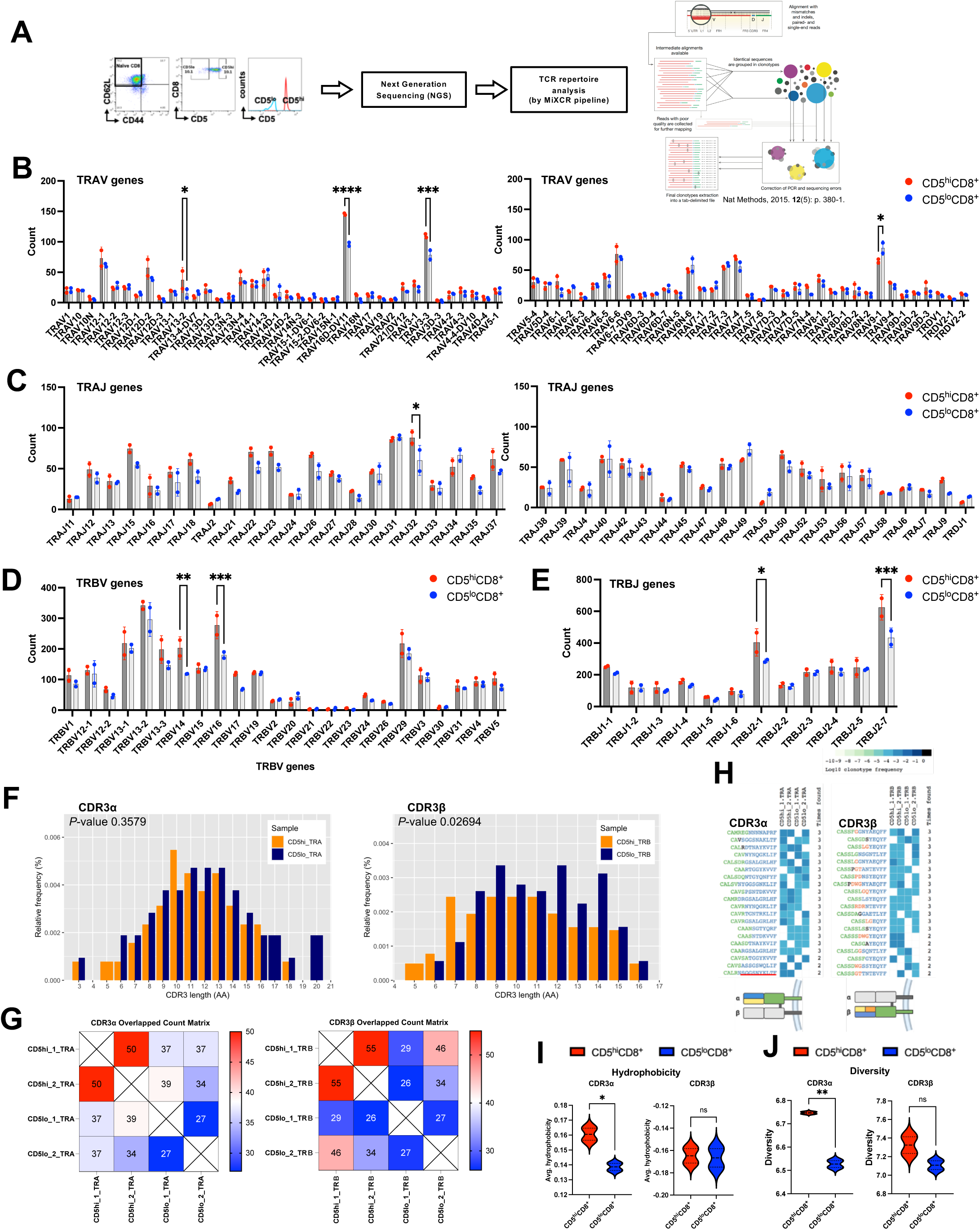
Examination of the CD5^hi^ population reveals TCR repertoires linked to autoimmune disorders. (**A**) Comparison of TCR repertoire differences between CD5^hi^CD8^+^ and CD5^lo^CD8^+^ T cell populations conducted using bulk RNA sequencing, followed by TCR repertoire analysis using the MiXCR pipeline. Naïve CD5^hi^CD8^+^ and CD5^lo^CD8^+^ cells were sorted from pooled spleens and PLNs of 6–8-week-old normoglycemic female NOD mice, following a similar procedure as outlined in Figure 2A, with 1.5 x 10^6^ cells per group. The TCR repertoire of naïve CD5^hi^CD8^+^ T cells exhibited shorter CDR3 lengths, higher clone overlap values within group comparisons, pathogenic-related CDR3 motifs and greater hydrophobicity and diversity in the TRA chain compared to CD8^+^CD5^lo^ cells. (**B**-**E**) Comparison of the TRAV gene segment usage (**B**), TRAJ gene segment usage (**C**), TRBV gene segment usage (**D**) and TRBJ gene segment usage (**E**) within TRA variable and TRB variable genes between CD5^hi^CD8^+^ (red) and CD5^lo^CD8^+^ (blue) cells. (**F**-**J**) Comparison of CDR3α (left) or CDR3β (right) length in amino acids (**F**), CDR3 clonal overlap by overlapped count matrix (**G**), top 20 CDR3 motifs (**H**), average hydrophobicity score (**I**) and diversity (**J**) of CDR3α (left) and CDR3β (right) between CD5^hi^CD8^+^ and CD5^lo^CD8^+^ T cells. In (**G**), the number in each cell indicates the count of shared clonotypes between sample pairs; the detailed method for constructing the Overlapped Count Matrix is provided in the Supplementary methods section. (**H**) Occurrence frequencies of the top 20 CDR3 motifs from mapped- CDR3α (left) and CDR3β (right) sequencings in CD5^hi^CD8^+^ and CD5^lo^CD8^+^ cells, with the high-occurrence rate of the CDR3α motif “SGGSNYKLTF” (underlined in red) specific to T1D-related IGRP peptide in the CD8^+^CD5^hi^ population. (**I**) Average hydrophobicity score defined by summing up the hydrophobicity score in each amino acid in the CDR3 motif and dividing by the CDR3 length, then averaging by total counts in each group. Diversity definition in (**J**) is shown in Materials and Methods section Clonotype repertoire metrics formulas. In (**B**-**E**), each point represents 10 mice, with two points for each group, totaling n = 20 mice (**F**-**J**) for all CD5^hi^ or CD5^lo^ samples. \**P* < 0.05, \*\**P* < 0.01, \*\*\**P* < 0.001, by two-way ANOVA (**B**-**E**), by Welch two sample t-test (**F**), and unpaired, two-tailed t test (**I**) and (**J**).

## Discussion

In our study, CD5^hi^CD8^+^ T cells, with heightened basal signals and increased self-ligand reactivity, show enhanced avidity for the autoantigen IGRP (Figure 8). These cells transition quickly into effector/memory cells upon encountering autoantigens, expressing markers like CD69 and CD44. Activation via CD122 and CD127 promotes cytokine secretion, contributing to inflammation and β cell destruction. The upregulation of CD5 in autoreactive CD8^+^ T cells within the PLNs may reflect ongoing antigen encounter and TCR engagement with self-peptides presented by dendritic cells or β cell-derived antigen-presenting cells. This localized antigen stimulation may enhance CD5 expression dynamically, as observed in previous studies where TCR self-reactivity correlates with CD5 levels in both naïve and memory T cells. Additionally, the increased activation marker expression (e.g., CD69 and CD44) in CD5^hi^CD8^+^ T cells suggests that these cells are actively engaging in self-antigen-driven priming, potentially accelerating their transition to an effector-like phenotype. Future studies employing *in vivo* antigen tracking or CD5 fate-mapping approaches could provide deeper insights into whether CD5^hi^ cells within the PLNs are newly activated by antigen encounter or preconditioned for heightened self-reactivity due to thymic selection. Moreover, further investigation is needed to understand the role of these memory-like CD8^+^ T cells in autoreactivity. The loss of Pep-conferred protection in high CD5- expressing NOD8.3 mice highlights the significance of elevated TCR basal signals from positive selection in autoimmune responses, suggesting potential targets for treating T1D and related disorders.

**Figure 8.**
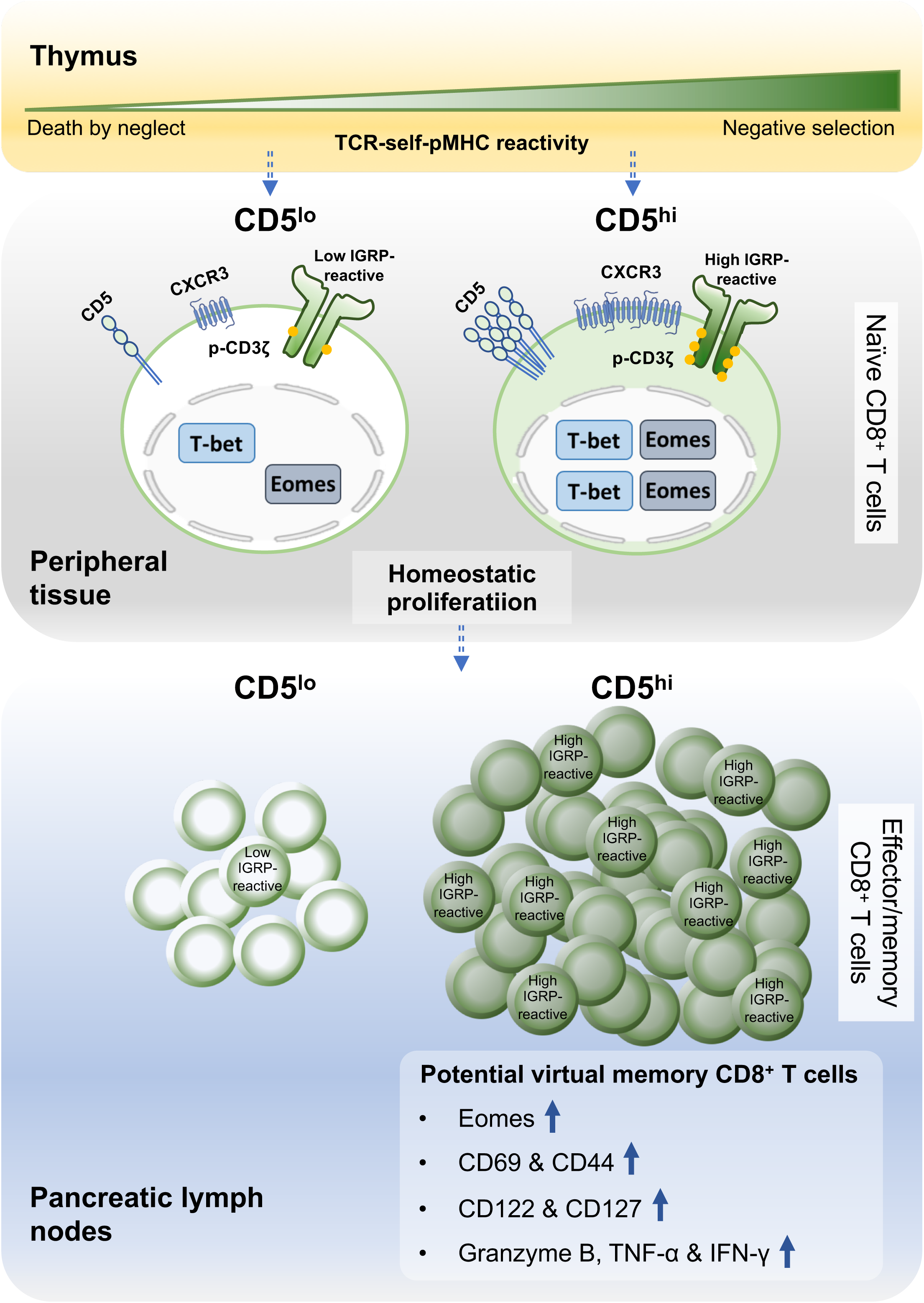
Thymocyte self-reactivity impacts the effector functions and memory cell formation of auto-reactive T cells in the T1D model NOD mice.

CD5 expression in DN thymocytes reflects the strength of pre-TCR signaling during early selection, where successful TCRβ rearrangement drives proliferation and differentiation into the DP stage. The phosphorylation of CD3ζ (pCD3ζ) at this stage indicates activation of Lck/ZAP-70 signaling, essential for lineage commitment and self-reactivity tuning. As addressed in a previous study, this early self-MHC interaction calibrates activation thresholds, ensuring appropriate TCR sensitivity while preventing excessive autoreactivity (61). The high expression of CD5 in DP TCRβ⁻CD69⁻ thymocytes from 8.3-TCR NOD mice may indicate early self-reactivity, even before overt TCR signaling. This raises the possibility that low-level self-pMHC recognition occurs at this stage, priming these cells for selection. To address this hypothesis, future experiments will assess TCR expression and activation states at this intermediate developmental phase, helping to clarify the role of CD5 in shaping the autoreactive potential of these cells. While bulk TCR-seq provided valuable insights into TCR repertoire differences between CD5^hi^ and CD5^lo^CD8^+^ T cells, it lacks the resolution to assess clonotype-specific self-reactivity. Single-cell TCR-seq would allow a more precise evaluation of individual T cell reactivity and clonal expansion, helping to determine whether CD5^hi^CD8^+^ cells preferentially retain higher-affinity self-reactive TCRs. Future studies will incorporate this approach to better define the relationship between CD5 expression and antigen specificity.

In contrast to previous studies that compared TCR signaling in NOD and wild-type B6 mice (11), we investigate how thymic self-recognition impacts TCR signal strength and CD8^+^ T cell pathogenicity in NOD mice. Our research also differs from studies focusing on the CD5^lo^CD4^+^ T-cell population (12), which resembles central memory T cells and is linked to spontaneous autoimmunity in NOD mice. Instead, we find distinct TCR signal strength variations between CD5^hi^ and CD5^lo^ cells, with CD5^hi^CD8^+^ cells showing enhanced activation, proliferation and disease transfer potential. This discrepancy may arise from the higher CD5-expressing distribution in the naïve CD4^+^ T cells compared to the naïve CD8^+^ T cells, suggesting distinct TCR signaling sensitivity when stimulated and resulting in different functional activation thresholds between CD4^+^ and CD8^+^ T cells (62, 63). Our results reveal that CD5^hi^CD8^+^ T cells exhibit elevated CD25 and CD122 expression compared to their CD5^lo^CD8^+^ counterparts, indicating increased sensitivity to IL-2 stimulation. While exposure to IL-2 may enhance cell expansion in CD8^+^ cells, it can lead to overstimulation in CD4^+^ T cells, resulting in activation-induced cell death. Surprisingly, clinical cases of type 1 diabetes reflect similar findings to our animal mouse model study. Children with T1D exhibit higher CD5 expression in CD8^+^ T cells compared to healthy controls (64). These observations underscore the complexity of immune responses in autoimmune diseases and warrant further exploration to unravel the underlying mechanisms and hold promise for therapeutic interventions in autoimmune diseases.

Nevertheless, the role of CD5 itself in shaping the distinctive functions of CD5^hi^ and CD5^lo^ cells remains largely unclear (65, 66). A previous research has indicated that CD5 serves as a scaffold protein, retaining the NF-κB inhibitor IκBα in the cytosol to prevent its inhibition of NF- κB signaling (67), though studies of Cd5^−/−^ mice do not appear to support this concept (68). This mechanism confers survival advantages to CD5^hi^ T cells. This observation may help elucidate the seemingly paradoxical result that CD5’s primary function is to attenuate TCR-proximal signaling; however interestingly, antigen-specific T cells with higher initial CD5 expression exhibit enhanced persistence as effector/memory cells following a peripheral challenge. Whether CD5^hi^CD8^+^ T cells execute potent autoimmune responses through the modulation of NF-κB-dependent signaling needs further investigation.

Exploring how TCR composition contributes to the distinct responses between CD5^hi^ and CD5^lo^ cells is beyond the scope of our study. Nonetheless, prior research, which involved screening autoimmune disease databases such as arthritis, diabetes, etc., has revealed a trend in specific TCR repertoire usage characterized by germline-encoded TCRs with fewer nucleotide insertions and shorter CDR3 length (60). It is consistent with our results, demonstrating that CD5^hi^ T cells, skewed toward shorter CDR3 lengths in the TCR repertoire, have a higher potential for autoreactivity. Consequently, the high self-pMHC reactivity of T cells provides a stronger TCR basal signal, potentially priming T cells with specific TCR compositions for robust responses to foreign antigens in the future. From an evolutionary perspective, this TCR design may have initially conferred an advantage by enhancing immune responses against pathogen infections. However, it might have come at the cost of an increased risk of autoimmune diseases. A previous study explored the TCR repertoire of CD8^+^ T cells recognizing the islet-specific peptide IGRP_206– 214_ in both grafted and endogenous islet infiltrates, revealing a shared repertoire favoring specific gene segments and limited diversity and clonotypic expansion within IGRP_206–214_-specific CD8^+^ T cells in NOD mice (58). Another study of effector/memory CD4^+^ and CD8^+^ T cells involved in autoimmune rejection of islet grafts in diabetic NOD mice demonstrated a comparable pattern of TCR repertoire diversity (69). Our analysis of TCR gene usage in CD5^hi^CD8^+^ and CD5^lo^CD8^+^ T cells also revealed distinct patterns, suggesting potential functional differences between these subsets. The identified differences included increased usage of TRAV13-2, TRAV16D-DV11, TRAV3-3, TRAJ32, TRBV14, TRBV16, TRBJ2-1, and TRBJ2-7 in CD5^hi^ compared to CD5^lo^ cells, while TRAV9-1 was reduced in CD5^hi^ cells. Notably, previous studies identified specific gene segments associated with autoimmune diseases like Sjögren’s syndrome (TRAV13-2) (70), inflammatory bowel disease, ulcerative colitis (TRBJ2-1 and TRBJ2-7) (71, 72), and type 1 diabetes (TRBV14 and TRBV16) (73), underscoring the importance of understanding T-cell receptor dynamics in autoimmune pathogenesis across various disorders.

Several studies have highlighted the impact of TCR-self-pMHC interactions during positive selection on signaling molecules’ localization, such as p-Erk, particularly in B cells and CD4^+^ T cells (4, 74, 75). Similarly, our CD5^hi^CD8^+^ T cells may consistently receive stronger signals, leading to better synchronization and activation of Erk at the cell membrane. However, despite using PMA/Iono to bypass proximal TCR signaling and elicit downstream responses, differences in T cell proliferation response persisted between CD5^hi^ and CD5^lo^ or between NOD and NOD8.3 CD8^+^ T cells. This suggests that TCR-self-peptide interaction influences T cells beyond just proximal TCR signaling intensity. There may be pre-existing transcriptional differences influenced by genes involved in T-cell receptor signaling established during thymic development among naïve NOD CD8^+^ T cells, affecting their effector lineage fate (76). Additionally, in a previous experimental autoimmune diabetes model, OVA antigen-specific CD8^+^ T cells with a virtual memory phenotype and high CD5 expression showed a low capacity to induce diabetes (77), contrasting with the pathogenic features of our CD5^hi^CD8^+^ cells. This suggests that naïve CD5^hi^CD8^+^ T cells in NOD mice may only partially resemble virtual memory T cells, posing intriguing questions requiring further investigation.

## Materials and methods

### Mice

The mice utilized in this study included NOD/Sytwu, NOD8.3, and NOD.BDC2.5 TCR transgenic mice, alongside Ptpn22-transgenic NOD mice, designated dLck–Ptpn22 C (dLPC/NOD) and dLck–Ptpn22 E (dLPE/NOD), previously generated by cloning Ptpn22 from NOD mouse splenocytes and microinjecting it into single-cell NOD embryos. The resulting dLck–Ptpn22 transgenic mice were genotyped by PCR and evaluated by Southern blot analysis for confirmation. Western blot analysis revealed the expression of transgenic Pep in thymocytes and splenic T cells of dLPC and dLPE mice, with a stepwise increase in Pep protein levels from nontransgenic littermates, dLPC/NOD to dLPE/NOD mice (27). The transgenic mice dLPC/NOD, dLPE/NOD, dLPC/NOD8.3 and dLPE/NOD8.3 used in the study were hemizygous for the Ptpn22 transgene (Supplementary Figure 7). dLPC/NOD8.3 and dLPE/NOD8.3 were established by mating dLPC/NOD or dLPE/NOD mice with NOD8.3 mice. NOD/Sytwu, NOD8.3 and NOD.BDC2.5 TCR transgenic mice were originally obtained from The Jackson Laboratory, while NOD Rag1^-/-^ mice were procured from the National Laboratory Animal Center. The study adhered to institutional guidelines and was approved by the National Defense Medical Center Institutional Animal Care and Use Committee.

### Sex as a biological variable

Our study exclusively examined female mice because the disease modeled is relevant in females.

### Statistics

The log-rank (Mantel–Cox) test was employed for survival curve comparisons. Two-group comparisons utilized a 2-tailed Student’s unpaired t-test, two-sample comparison utilized Welch two sample t-test, and multigroup comparisons utilized one-way ANOVA with Tukey’s post-test. A significance level of *P* < 0.05 was considered significant.

### Study approval

All protocols involving live animals adhered to institutional guidelines and received approval from the Institutional Animal Care and Use Committee at the National Defense Medical Center (Taipei, Taiwan).

Additional methods can be found in the Supplementary Material.

*Supplementary material.* Supplementary material provided with this study includes Supplementary Figures 1-7, Supplementary Tables 1-6 and Supplementary methods.

## Data availability

The TCR repertoires of CD5^hi^ and CD5^lo^, along with the data analyzed using R and the corresponding codes, are available in the following GitHub repository: https://github.com/Chia-Lo/TCR-signal-strength-modulates-antigen-specific-CD8-T-cell-pathogenicity-in-non-obese-diabetic-mice.git

## Author contributions

CLH and LTY performed experiments and analyzed data. CLH, LTY, YWL and JLD performed experiments. LTY gave advice. CLH and HKS wrote the manuscript.

## Acknowledgments

This work was supported by the Ministry of Science and Technology of the Republic of China (MOST 109-2320-B-400-018-MY3, MOST 110-2320-B-400-011-MY3), National Science and Technology Council (NSTC 112-2320-B-400-026-MY3) and Tri-Service General Hospital (TSGH-C02-112029, TSGH-C03-113037, VTA112-T-1-1, VTA113-T-1-1). The authors acknowledge the technical services provided by Instrument Center of National Defense Medical Center.

## Abbreviations

dLPC/NOD: *Lck* distal promoter– *Ptpn22* C line in NOD background
dLPE/NOD: *Lck* distal promoter– *Ptpn22* E line in NOD background
NOD: non-obese diabetic.

## Supplementary Figures

**Supplementary Figure 1.**
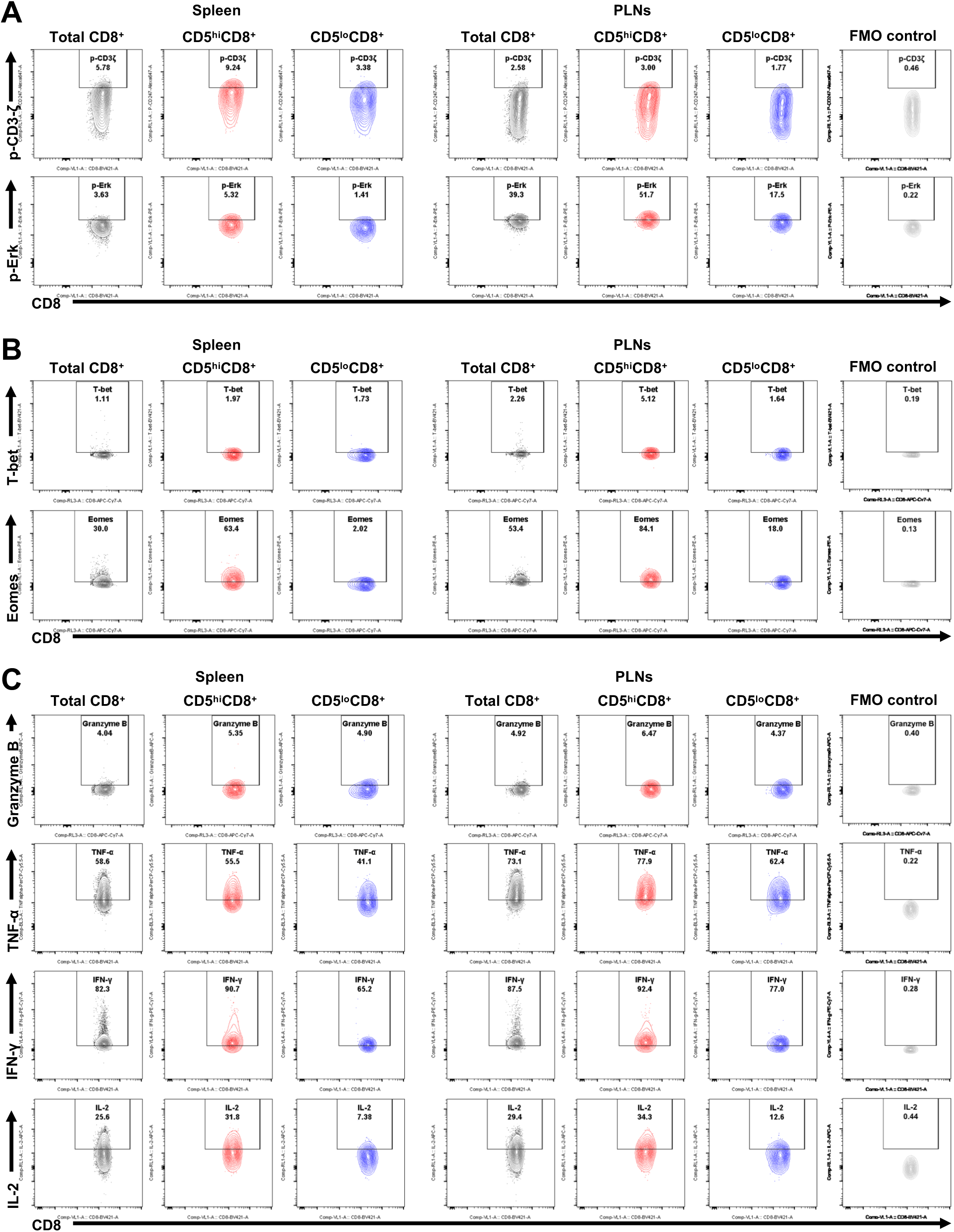
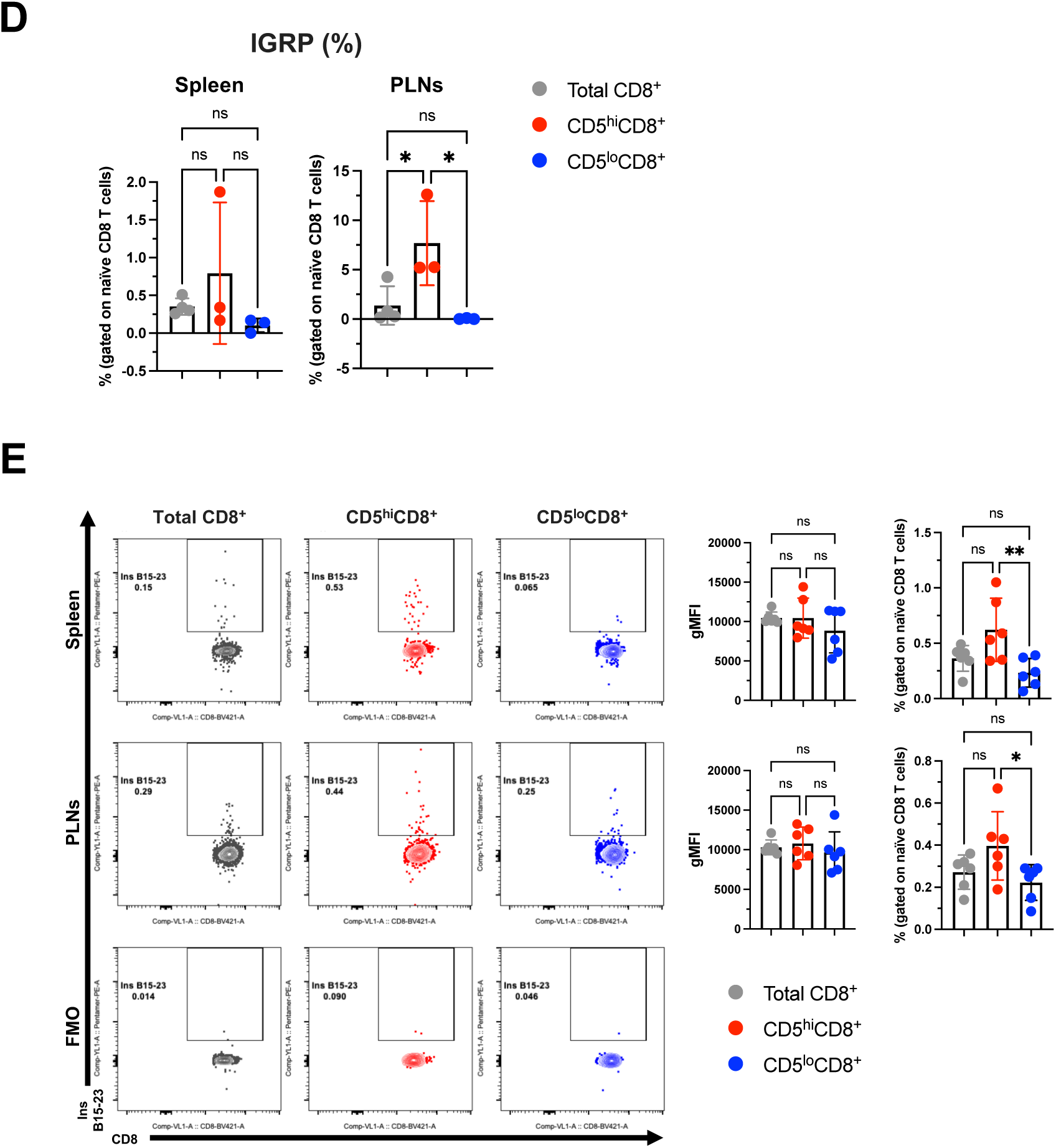
CD5^hi^CD8^+^ T cells display enhanced TCR signaling, effector profiles and autoreactive potential in NOD mice. Representative FLOW analysis plots show a comparison of positive expression percentages in (**A**) proximal TCR signaling molecules p-CD3ζ and p-Erk, (**B**) transcription factors T-bet and Eomes, and (**C**) cytokines Granzyme B, TNF-α, IFN-γ and IL-2 within total naïve CD8^+^ (grey), CD5^hi^CD8^+^ (red), CD5^lo^CD8^+^ (blue) cells, and fluorescence minus one (FMO) controls (light grey) from spleen (left) and PLNs (right). (**D**) Corresponding percentages of IGRP-tetramer positive CD8^+^ T cells among total naïve CD8^+^ T cells, CD5^hi^CD8^+^ and CD5^lo^CD8^+^ populations from spleen and PLNs. (**E**) Representative flow cytometry plots (left) and corresponding gMFI and proportions (right) of Ins B15-23-pentamer positive CD8^+^ T cells compared among total naïve CD8^+^ T cells, CD5^hi^CD8^+^ and CD5^lo^CD8^+^ populations from spleen and PLNs. In (**D**) and (**E**) all panels, total naïve CD8^+^ T cells, CD5^hi^CD8^+^ and CD5^lo^CD8^+^ are colored grey, red and blue, respectively. Data represent mean ± SD. Sample size was n = 3–5 per group. \**P* < 0.05, \*\**P* < 0.01, by one-way ANOVA.

**Supplementary Figure 2.**
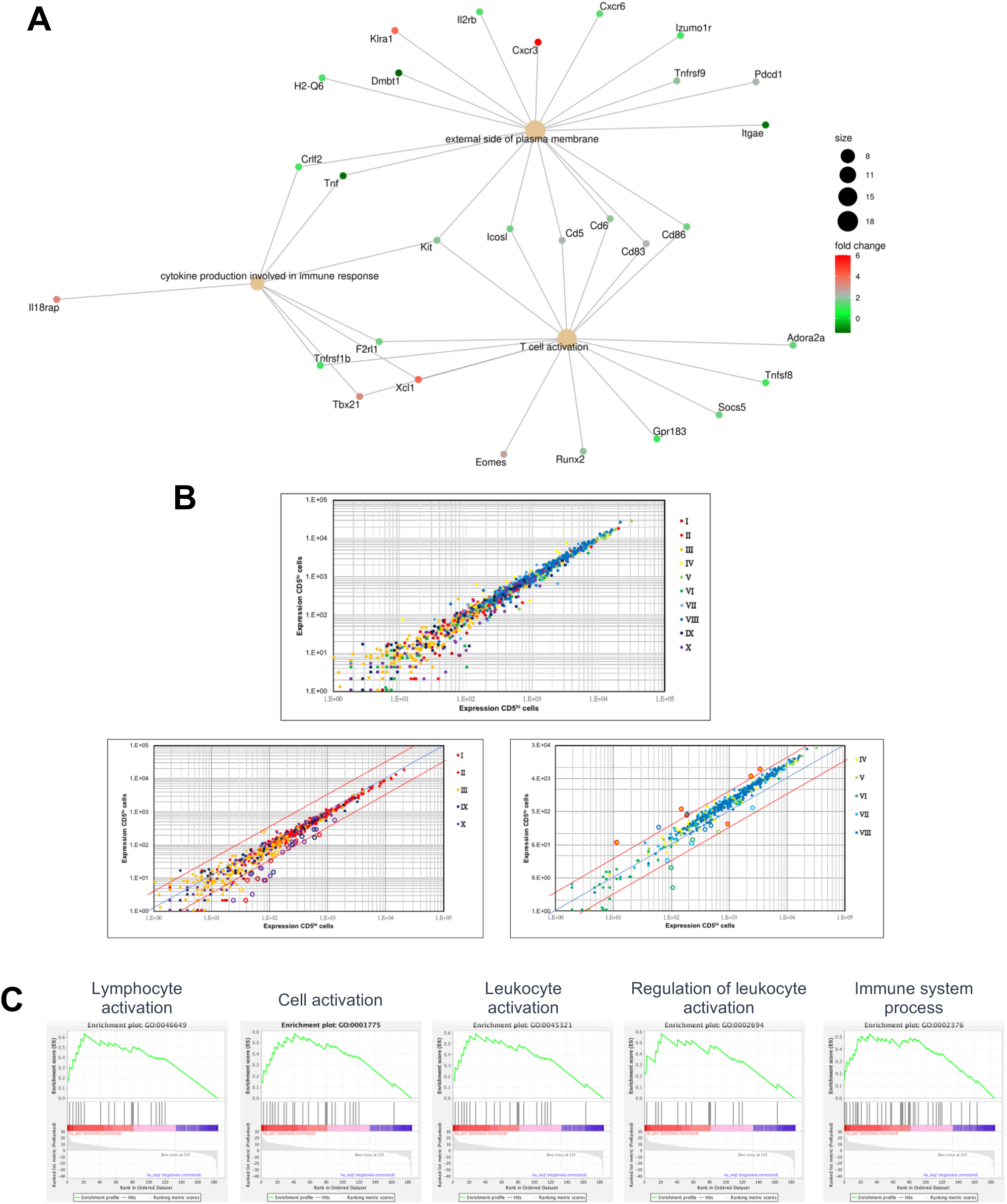
Gene expression patterns and enrichment are comparatively analyzed between naïve CD5^hi^ and CD5^lo^CD8^+^ T cells. (**A**) Gene-concept network of CD5^hi^ versus CD5^lo^ DEGs across key immune process GO terms. This interaction picture displays a gene- concept network highlighting genes associated with the top three GO terms: “external side of plasma membrane” (GO:0009897), “T cell activation” (GO:0042110), and “cytokine production involved in immune response” (GO:0002367) in CD5^hi^ versus CD5^lo^ DEGs. Circle size represents the gene counts, while the color spectrum, ranging from green to red, indicates the fold change in gene expression, spanning from less than 0 to a 6-fold change. (**B**) Comparison of gene expression patterns of CD5^hi^CD8^+^ (x-axis) versus CD5^lo^CD8^+^ (y-axis) T cells, using core gene signatures from clusters I to X (top panel). The two lower panels represent a similar comparison to the top panel but specifically focusing on the core gene signature differences in (left) clusters I, II, III, IX, and X, and (right) clusters IV to VIII between CD5^hi^ and CD5^lo^ T cells (lower panel); colors in plots (genes) match colors of clusters (key); red diagonal lines indicate a difference in expression of twofold; blue diagonal line indicates x = y. \**P* < 0.001 and \*\**P* < 0.00001 (t-test). Data are representative of two independent experiments. (**C**) The gene set enrichment score plot for biological process (BP) of GO terms (lymphocyte activation, cell activation, leukocyte activation, regulation of leukocyte activation, and immune system process). Higher scores indicate a higher degree of enrichment of these processes in naïve CD5^hi^CD8^+^ than in CD5^lo^CD8^+^ cells.

**Supplementary Figure 3.**
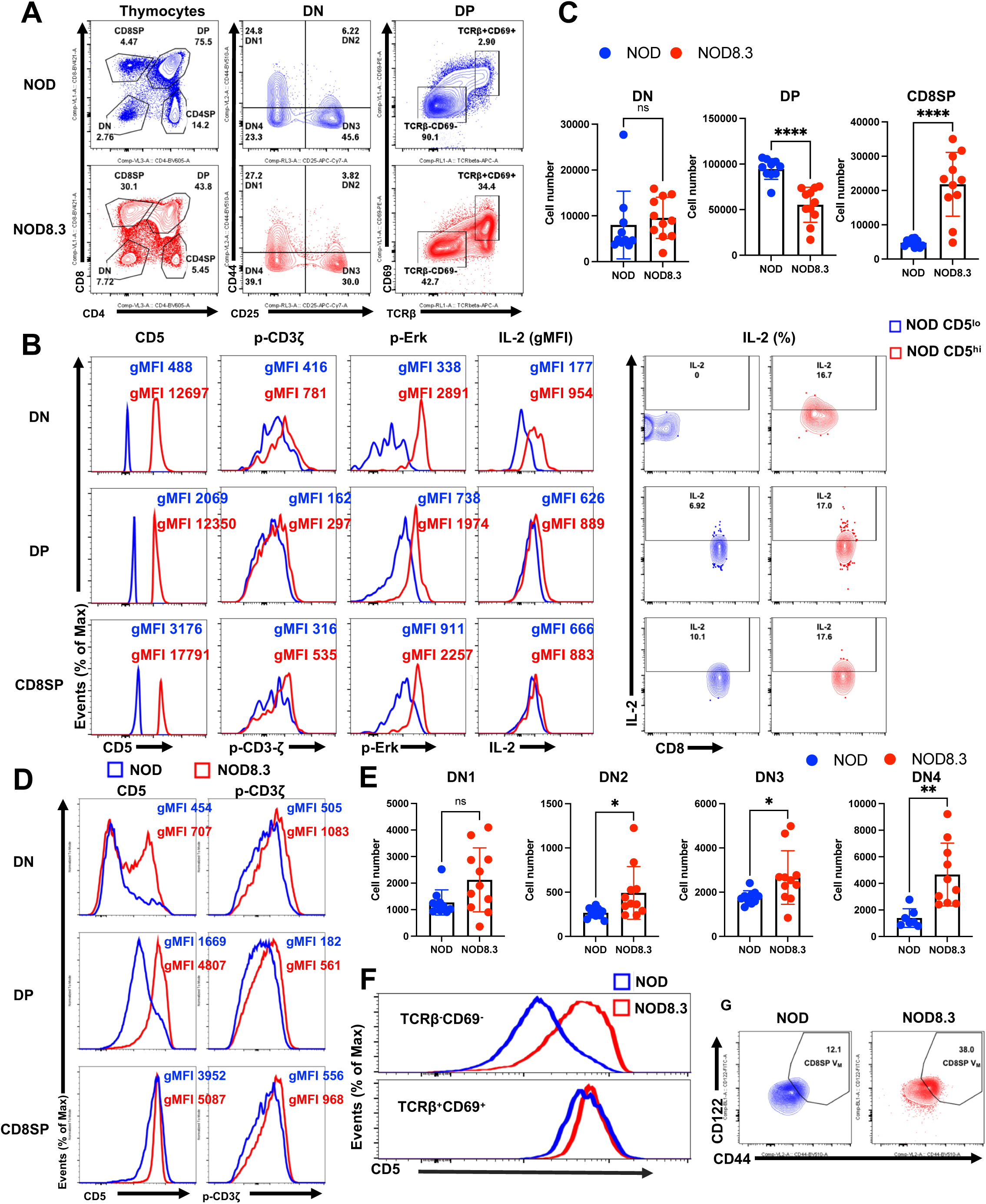
Thymocyte development and respective selection profiles are compared between NOD and NOD8.3 mice. (**A**) Representative FLOW dot plots showing thymocytes (CD4 and CD8), DN thymocytes (CD25 and CD44), pre-positive selection (TCRβ^-^CD69^-^), and post-positive selection (TCRβ^+^CD69^+^) stages in DP thymocytes from NOD and NOD 8.3 mice. (**B**) Representative gMFI histograms of CD5, p- CD3ζ, p-Erk and IL-2 levels (left) and the IL-2 gating representative plot (right) compared between the lower 5% and upper 5% CD5 expression in thymocytes of normoglycemia female NOD mice aged 6 to 8 weeks old across DN, DP and CD8SP stages. (**C**) Cell numbers of thymocytes from NOD and NOD8.3 mice at the DN, DP and CD8SP stages. (**D**) Representative CD5 and p-CD3ζ gMFI histograms in thymocytes across DN, DP and CD8SP stages from NOD and NOD8.3 mice. (**E**) Cell numbers of thymocytes at the indicated DN stages (DN1, DN2, DN3 and DN4) from NOD and NOD8.3 mice. DN stages are defined as presented in Supplementary Figure 3A, middle panels. (**F**) CD5 gMFI distribution at the pre-positive selection (upper; TCRβ^-^CD69^-^) or post-positive selection (lower; TCRβ^+^CD69^+^) stage in DP thymocytes from NOD and NOD8.3 mice. (**G**) Representative FLOW dot plots for virtual memory (Vmemory) as shown in Figure 5H in NOD or NOD8.3 mice. In all panels from (**A**) to (**G**), NOD CD5^lo^ cells or NOD mice are represented in blue, and NOD CD5^hi^ cells or NOD8.3 mice are represented in red. Data represent mean ± SD. The data presented in (**C**) and (**E**) represent two experiments. The sample size was n = 10-11 (**C**) and n = 7-11 (**E**) per group. \**P* < 0.05, \*\**P* < 0.01, \*\*\*\**P* < 0.0001, by unpaired, two-tailed t test (**C**) and (**E**).

**Supplementary Figure 4.**
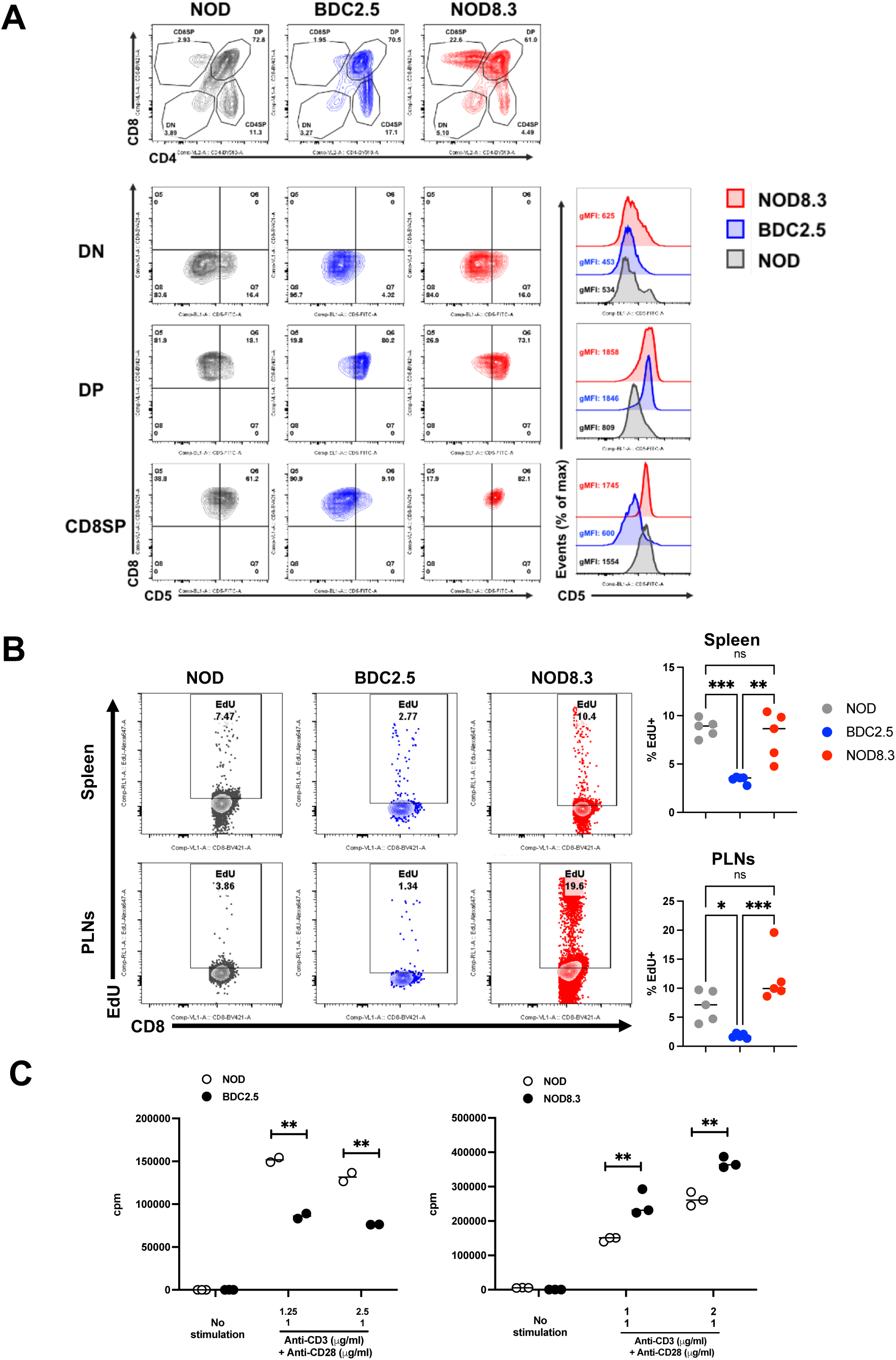
Thymocyte-stage self-reactivity correlates peripheral diabetogenic T cell proliferation profiles in NOD, BDC2.5 and NOD8.3 mice. (**A**) CD5 gMFI comparison in the DN, DP and CD8SP stages of thymocytes (upper panel) among NOD (grey), BDC2.5 (blue) and NOD8.3 (red) mice. The received TCR signal (dictated by CD5 level) during thymocyte selection dictates the peripheral cell proliferation level, shown in (**B** and **C**). (**B**) Thymidine analog Edu incorporation in CD8^+^ T cells from spleen (upper) or PLNs (lower) compared among NOD (grey), BDC2.5 (blue) and NOD8.3 (red) mice. Representative FLOW cytometry dot plots (left) and corresponding statistical results (right) are presented. (**C**) T cell proliferation level in BDC2.5 (left panel) and NOD8.3 (right panel) compared to NOD mice. Naïve T cells were isolated from 6-8-week-old normoglycemia female NOD, BDC2.5 and NOD8.3 mice and stimulated with the indicated amounts of anti-CD3 and anti-CD28 (1 μg/ml). Cell proliferation was measured by the incorporation of [methyl-3H] thymidine. Data represent mean ± SD. The data presented in (**B**) represent two experiments. The sample size was n = 5 (**B**) and n = 2-3 (**C**) per group. \**P* < 0.05, \*\**P* < 0.01, \*\*\**P* < 0.001, by one-way ANOVA (**B**) and unpaired, two-tailed t test (**C**).

**Supplementary Figure 5.**
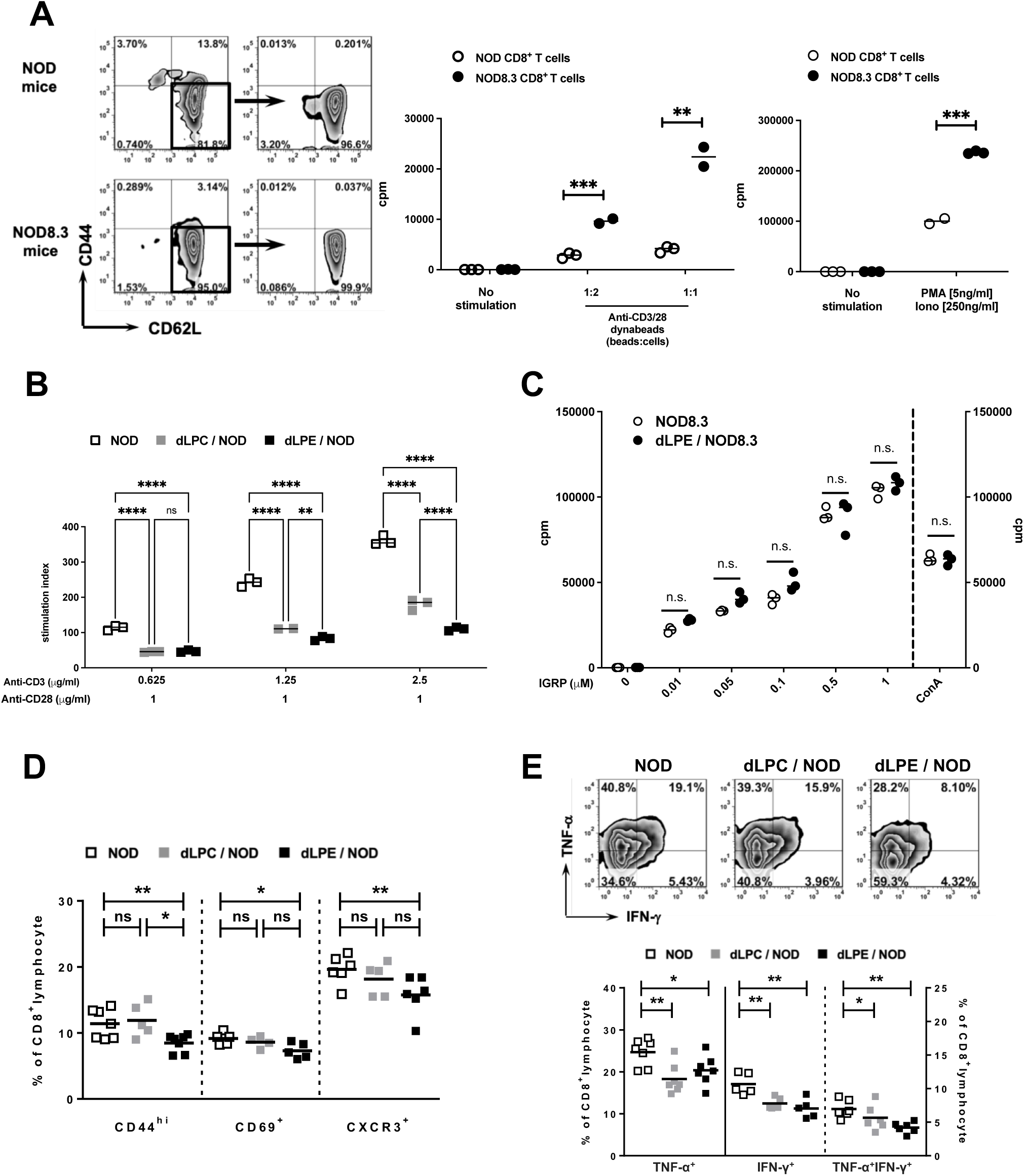
Immunomodulatory effects of transgenic Pep on effector T cell functions are analyzed. (**A**) Cell proliferation levels of naïve CD44^lo^CD62L^hi^CD8^+^ T cells (left) isolated from 8-10-week-old normoglycemia female NOD or NOD8.3 upon stimulation with anti- CD3/CD28 dynabeads (middle) or PMA/Ionomycin (right) treatment, measured by the incorporation of [methyl-3H] thymidine (cpm; counts per minute). (**B**) Cell proliferation levels of CD8^+^ T cells isolated from 8-10-week-old normoglycemia female NOD, dLPC/NOD and dLPE/NOD mice upon anti-CD3/CD28 stimulation. Purified CD8^+^ T cells were stimulated with the indicated amounts of anti-CD3 (0.625, 1.25 and 2.5 μg/ml, respectively) combined with anti-CD28 (1 μg/ml). Cell proliferation was measured by the incorporation of [methyl-3H] thymidine (shown by simulation index). (**C**) Proliferation levels of splenocytes isolated from 8-10-week-old normoglycemia female NOD8.3 and dLPE/NOD8.3 mice upon stimulation with the indicated amounts of IGRP_206-214_ peptides or ConA. Cell proliferation was measured by the incorporation of [methyl-3H] thymidine (cpm; counts per minute). (**D**) Percentages of effector/memory CD44^hi^-, CD69^+^-, and CXCR3^+^CD8^+^ T cells in PLNs of NOD, dLPC/NOD and dLPE/NOD mice. (**E**) Percentages of TNF-α^+^-, IFN-γ^+^-, and TNF-α^+^IFN-γ^+^CD8^+^ T cells in 8-10-week-old normoglycemia female NOD, dLPC/NOD and dLPE/NOD mice. Isolated cells were stimulated with anti-CD3/CD28 (1 μg/ml) for 48 hours. After stimulation, the percentages of cytokine-producing CD8^+^ T cells were measured by flow cytometry analysis. Notably, in (**B**), (**D**) and (**E**), a stepwise increase in transgenic Pep expression from nontransgenic NOD to dLPC/NOD and dLPE/NOD mice within the litters born at the same time was confirmed in our previous study (27). Data represent mean ± SD. The data presented in (**A**) to (**E**) represent two experiments. The sample size was n = 5-7 (**A**), n = 5-7 (**B**), n = 6 (**C**), n = 5-7 (**D**) and n = 5-7 (**E**) per group. \**P* < 0.05, \*\**P* < 0.01, \*\*\**P* < 0.001, by unpaired, two-tailed t test (**A**) and (**C**) and one-way ANOVA (**B**, **D** and **E**).

**Supplementary Figure 6.**
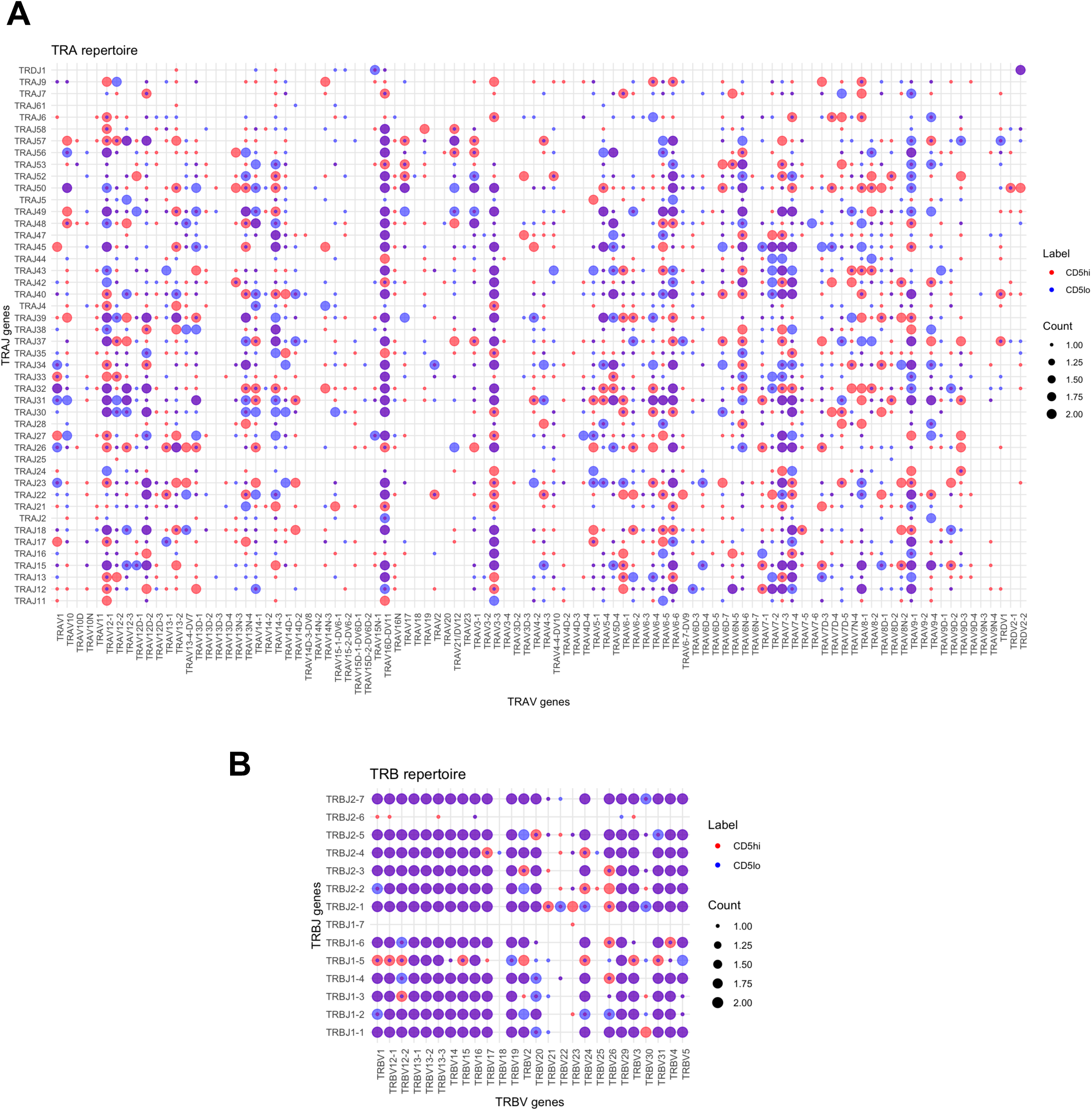
The pairing of VJ gene segments in the TCR repertoire is compared between CD5^hi^CD8^+^ and CD5^lo^CD8^+^ cells. **(A)** TRA variable (TRAV) and **(B)** TRB variable (TRBV) genes in CD5^hi^ (red dot) and CD5^lo^ (blue dot) groups were examined. Notably, if the same count of a specific VJ gene segment usage occurred in CD5^hi^ and CD5^lo^ cells, the outcome of overlapping dot color would be purple. n = 20 mice for each CD5^hi^ and CD5^lo^ group.

**Supplementary Figure 7.**
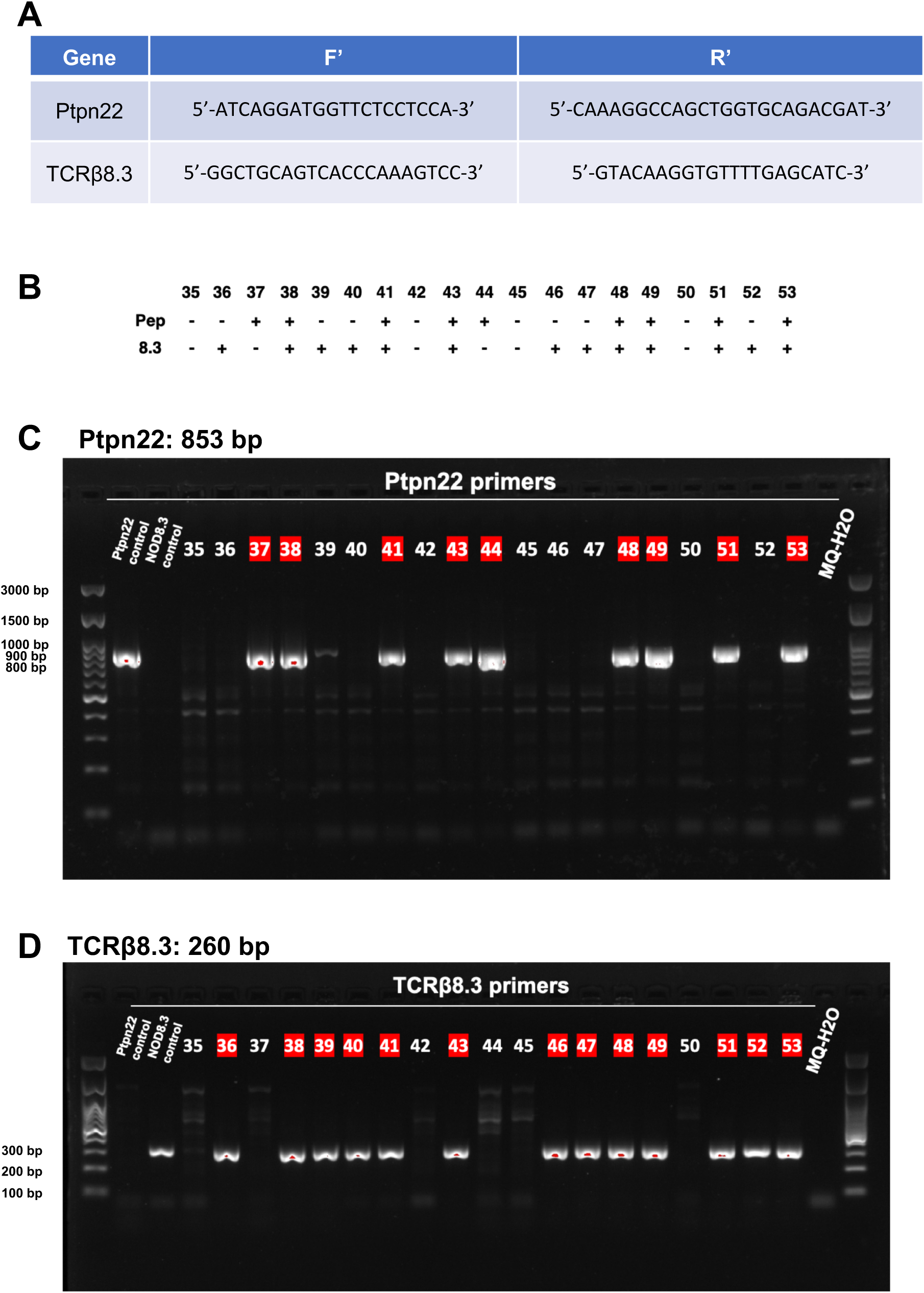
The genotyping results of NOD, dLPC/NOD, NOD8.3 and dLPC/NOD8.3 mice using the indicated F’ and R’ primers are shown. **(A)** The Ptpn22 F’ and R’ primers used for genotyping were designed in our previous study to evaluate the existence of the transgene Ptpn22 (Ptpn22: 853 bp) (27). The primer sequence for TCRβ8.3 can be referred to (Verdaguer et al., 1997) (TCRβ8.3: 260 bp). **(B-D)** Genotyping results for transgenic Ptpn22 and TCRβ8.3 expression in pups #35 to #53 from male NOD8.3 crossed with female dLPC/NOD mice are presented.

## References

1. Gascoigne NR, Rybakin V, Acuto O, and Brzostek J. TCR Signal Strength and T Cell Development. Annu Rev Cell Dev Biol. 2016;32:327–48.

2. Persaud SP, Parker CR, Lo WL, Weber KS, and Allen PM. Intrinsic CD4+ T cell sensitivity and response to a pathogen are set and sustained by avidity for thymic and peripheral complexes of self peptide and MHC. Nat Immunol. 2014;15(3):266–74.

3. Fulton RB, Hamilton SE, Xing Y, Best JA, Goldrath AW, Hogquist KA, et al. The TCR’s sensitivity to self peptide-MHC dictates the ability of naive CD8(+) T cells to respond to foreign antigens. Nat Immunol. 2015;16(1):107–17.

4. Myers DR, Zikherman J, and Roose JP. Tonic Signals: Why Do Lymphocytes Bother? Trends Immunol. 2017;38(11):844–57.

5. Mandl JN, Monteiro JP, Vrisekoop N, and Germain RN. T cell-positive selection uses self- ligand binding strength to optimize repertoire recognition of foreign antigens. Immunity. 2013;38(2):263–74.

6. Deshpande NR, Parrish HL, and Kuhns MS. Self-recognition drives the preferential accumulation of promiscuous CD4(+) T-cells in aged mice. Elife. 2015;4:e05949.

7. Sadigh S, Scott BB, Mageed RA, Malcolm A, Andrew EM, and Maini RN. Identification of hybridomas derived from mouse CD5+ B lymphocytes by fluorescent staining for cytoplasmic CD5 expression. Immunology. 1994;81(4):558–63.

8. Tarakhovsky A, Kanner SB, Hombach J, Ledbetter JA, Muller W, Killeen N, et al. A role for CD5 in TCR-mediated signal transduction and thymocyte selection. Science. 1995;269(5223):535-7.

9. Perez-Villar JJ, Whitney GS, Bowen MA, Hewgill DH, Aruffo AA, and Kanner SB. CD5 negatively regulates the T-cell antigen receptor signal transduction pathway: involvement of SH2-containing phosphotyrosine phosphatase SHP-1. Mol Cell Biol. 1999;19(4):2903–12.

10. Azzam HS, Grinberg A, Lui K, Shen H, Shores EW, and Love PE. CD5 expression is developmentally regulated by T cell receptor (TCR) signals and TCR avidity. J Exp Med. 1998;188(12):2301–11.

11. Dong M, Audiger C, Adegoke A, Lebel ME, Valbon SF, Anderson CC, et al. CD5 levels reveal distinct basal T-cell receptor signals in T cells from non-obese diabetic mice. Immunol Cell Biol. 2021.

12. Kong Y, Jing Y, Allard D, Scavuzzo MA, Sprouse ML, Borowiak M, et al. A dormant T- cell population with autoimmune potential exhibits low self-reactivity and infiltrates islets in type 1 diabetes. Eur J Immunol. 2022.

13. Lutes LK, Steier Z, McIntyre LL, Pandey S, Kaminski J, Hoover AR, et al. T cell self- reactivity during thymic development dictates the timing of positive selection. Elife. 2021;10.

14. Han B, Serra P, Yamanouchi J, Amrani A, Elliott JF, Dickie P, et al. Developmental control of CD8 T cell-avidity maturation in autoimmune diabetes. J Clin Invest. 2005;115(7):1879–87.

15. Amrani A, Verdaguer J, Serra P, Tafuro S, Tan R, and Santamaria P. Progression of autoimmune diabetes driven by avidity maturation of a T-cell population. Nature. 2000;406(6797):739–42.

16. Mingueneau M, Jiang W, Feuerer M, Mathis D, and Benoist C. Thymic negative selection is functional in NOD mice. J Exp Med. 2012;209(3):623–37.

17. Eisenbarth GS. Type I diabetes mellitus. A chronic autoimmune disease. N Engl J Med. 1986;314(21):1360–8.

18. Ziegler AG, and Nepom GT. Prediction and pathogenesis in type 1 diabetes. Immunity. 2010;32(4):468–78.

19. Herold KC, Vignali DA, Cooke A, and Bluestone JA. Type 1 diabetes: translating mechanistic observations into effective clinical outcomes. Nat Rev Immunol. 2013;13(4):243–56.

20. Pearson JA, Wong FS, and Wen L. The importance of the Non Obese Diabetic (NOD) mouse model in autoimmune diabetes. J Autoimmun. 2016;66:76–88.

21. Anderson MS, and Bluestone JA. The NOD mouse: a model of immune dysregulation. Annu Rev Immunol. 2005;23:447–85.

22. Fousteri G, Liossis SN, and Battaglia M. Roles of the protein tyrosine phosphatase PTPN22 in immunity and autoimmunity. Clin Immunol. 2013;149(3):556–65.

23. Ivashkiv LB. PTPN22 in autoimmunity: different cell and different way. Immunity. 2013;39(1):91–3.

24. Bottini N, and Peterson EJ. Tyrosine phosphatase PTPN22: multifunctional regulator of immune signaling, development, and disease. Annu Rev Immunol. 2014;32:83–119.

25. Verdaguer J, Yoon JW, Anderson B, Averill N, Utsugi T, Park BJ, et al. Acceleration of spontaneous diabetes in TCR-beta-transgenic nonobese diabetic mice by beta-cell cytotoxic CD8+ T cells expressing identical endogenous TCR-alpha chains. J Immunol. 1996;157(10):4726–35.

26. Verdaguer J, Schmidt D, Amrani A, Anderson B, Averill N, and Santamaria P. Spontaneous autoimmune diabetes in monoclonal T cell nonobese diabetic mice. J Exp Med. 1997;186(10):1663–76.

27. Yeh LT, Miaw SC, Lin MH, Chou FC, Shieh SJ, Chuang YP, et al. Different modulation of Ptpn22 in effector and regulatory T cells leads to attenuation of autoimmune diabetes in transgenic nonobese diabetic mice. J Immunol. 2013;191(2):594–607.

28. Kaech SM, and Cui W. Transcriptional control of effector and memory CD8+ T cell differentiation. Nat Rev Immunol. 2012;12(11):749–61.

29. Pearce EL, Mullen AC, Martins GA, Krawczyk CM, Hutchins AS, Zediak VP, et al. Control of effector CD8+ T cell function by the transcription factor Eomesodermin. Science. 2003;302(5647):1041-3.

30. Intlekofer AM, Takemoto N, Wherry EJ, Longworth SA, Northrup JT, Palanivel VR, et al. Effector and memory CD8+ T cell fate coupled by T-bet and eomesodermin. Nat Immunol. 2005;6(12):1236–44.

31. Zhu Y, Ju S, Chen E, Dai S, Li C, Morel P, et al. T-bet and eomesodermin are required for T cell-mediated antitumor immune responses. J Immunol. 2010;185(6):3174–83.

32. Lord GM, Rao RM, Choe H, Sullivan BM, Lichtman AH, Luscinskas FW, et al. T-bet is required for optimal proinflammatory CD4+ T-cell trafficking. Blood. 2005;106(10):3432–9.

33. Banerjee A, Gordon SM, Intlekofer AM, Paley MA, Mooney EC, Lindsten T, et al. Cutting edge: The transcription factor eomesodermin enables CD8+ T cells to compete for the memory cell niche. J Immunol. 2010;185(9):4988–92.

34. Palmer MJ, Mahajan VS, Chen J, Irvine DJ, and Lauffenburger DA. Signaling thresholds govern heterogeneity in IL-7-receptor-mediated responses of naive CD8(+) T cells. Immunol Cell Biol. 2011;89(5):581–94.

35. Boyman O, Letourneau S, Krieg C, and Sprent J. Homeostatic proliferation and survival of naive and memory T cells. Eur J Immunol. 2009;39(8):2088–94.

36. Lenz DC, Kurz SK, Lemmens E, Schoenberger SP, Sprent J, Oldstone MB, et al. IL-7 regulates basal homeostatic proliferation of antiviral CD4+T cell memory. Proc Natl Acad Sci U S A. 2004;101(25):9357–62.

37. Rubinstein MP, Lind NA, Purton JF, Filippou P, Best JA, McGhee PA, et al. IL-7 and IL-15 differentially regulate CD8+ T-cell subsets during contraction of the immune response. Blood. 2008;112(9):3704–12.

38. Cibrian D, and Sanchez-Madrid F. CD69: from activation marker to metabolic gatekeeper. Eur J Immunol. 2017;47(6):946–53.

39. Rea IM, McNerlan SE, and Alexander HD. CD69, CD25, and HLA-DR activation antigen expression on CD3+ lymphocytes and relationship to serum TNF-alpha, IFN-gamma, and sIL-2R levels in aging. Exp Gerontol. 1999;34(1):79–93.

40. Ahn E, Araki K, Hashimoto M, Li W, Riley JL, Cheung J, et al. Role of PD-1 during effector CD8 T cell differentiation. Proc Natl Acad Sci U S A. 2018;115(18):4749–54.

41. Agata Y, Kawasaki A, Nishimura H, Ishida Y, Tsubata T, Yagita H, et al. Expression of the PD-1 antigen on the surface of stimulated mouse T and B lymphocytes. Int Immunol. 1996;8(5):765–72.

42. Herndler-Brandstetter D, Ishigame H, Shinnakasu R, Plajer V, Stecher C, Zhao J, et al. KLRG1(+) Effector CD8(+) T Cells Lose KLRG1, Differentiate into All Memory T Cell Lineages, and Convey Enhanced Protective Immunity. Immunity. 2018;48(4):716–29 e8.

43. Park JH, Adoro S, Lucas PJ, Sarafova SD, Alag AS, Doan LL, et al. ’Coreceptor tuning’: cytokine signals transcriptionally tailor CD8 coreceptor expression to the self-specificity of the TCR. Nat Immunol. 2007;8(10):1049–59.

44. Cho JH, Kim HO, Surh CD, and Sprent J. T cell receptor-dependent regulation of lipid rafts controls naive CD8+ T cell homeostasis. Immunity. 2010;32(2):214–26.

45. Lovering RC, Camon EB, Blake JA, and Diehl AD. Access to immunology through the Gene Ontology. Immunology. 2008;125(2):154–60.

46. Dorner BG, Dorner MB, Zhou X, Opitz C, Mora A, Guttler S, et al. Selective expression of the chemokine receptor XCR1 on cross-presenting dendritic cells determines cooperation with CD8+ T cells. Immunity. 2009;31(5):823–33.

47. Best JA, Blair DA, Knell J, Yang E, Mayya V, Doedens A, et al. Transcriptional insights into the CD8(+) T cell response to infection and memory T cell formation. Nat Immunol. 2013;14(4):404–12.

48. Godfrey DI, Kennedy J, Suda T, and Zlotnik A. A developmental pathway involving four phenotypically and functionally distinct subsets of CD3-CD4-CD8- triple-negative adult mouse thymocytes defined by CD44 and CD25 expression. J Immunol. 1993;150(10):4244–52.

49. Michie AM, and Zuniga-Pflucker JC. Regulation of thymocyte differentiation: pre-TCR signals and beta-selection. Semin Immunol. 2002;14(5):311–23.

50. Yamasaki S, and Saito T. Molecular basis for pre-TCR-mediated autonomous signaling. Trends Immunol. 2007;28(1):39–43.

51. Hoffman ES, Passoni L, Crompton T, Leu TM, Schatz DG, Koff A, et al. Productive T-cell receptor beta-chain gene rearrangement: coincident regulation of cell cycle and clonality during development in vivo. Genes Dev. 1996;10(8):948–62.

52. Fu G, Vallee S, Rybakin V, McGuire MV, Ampudia J, Brockmeyer C, et al. Themis controls thymocyte selection through regulation of T cell antigen receptor-mediated signaling. Nat Immunol. 2009;10(8):848–56.

53. Kim S, Park GY, Park JS, Park J, Hong H, and Lee Y. Regulation of positive and negative selection and TCR signaling during thymic T cell development by capicua. Elife. 2021;10.

54. Miller CH, Klawon DEJ, Zeng S, Lee V, Socci ND, and Savage PA. Eomes identifies thymic precursors of self-specific memory-phenotype CD8(+) T cells. Nat Immunol. 2020;21(5):567–77.

55. Hughes MM, Yassai M, Sedy JR, Wehrly TD, Huang CY, Kanagawa O, et al. T cell receptor CDR3 loop length repertoire is determined primarily by features of the V(D)J recombination reaction. Eur J Immunol. 2003;33(6):1568–75.

56. Gomez-Tourino I, Kamra Y, Baptista R, Lorenc A, and Peakman M. T cell receptor beta- chains display abnormal shortening and repertoire sharing in type 1 diabetes. Nat Commun. 2017;8(1):1792.

57. Zvyagin IV, Pogorelyy MV, Ivanova ME, Komech EA, Shugay M, Bolotin DA, et al. Distinctive properties of identical twins’ TCR repertoires revealed by high-throughput sequencing. Proc Natl Acad Sci U S A. 2014;111(16):5980–5.

58. Fuchs YF, Eugster A, Dietz S, Sebelefsky C, Kuhn D, Wilhelm C, et al. CD8(+) T cells specific for the islet autoantigen IGRP are restricted in their T cell receptor chain usage. Sci Rep. 2017;7:44661.

59. Stadinski BD, Shekhar K, Gomez-Tourino I, Jung J, Sasaki K, Sewell AK, et al. Hydrophobic CDR3 residues promote the development of self-reactive T cells. Nat Immunol. 2016;17(8):946–55.

60. Madi A, Shifrut E, Reich-Zeliger S, Gal H, Best K, Ndifon W, et al. T-cell receptor repertoires share a restricted set of public and abundant CDR3 sequences that are associated with self-related immunity. Genome Res. 2014;24(10):1603–12.

61. Huseby ES, and Teixeiro E. The perception and response of T cells to a changing environment are based on the law of initial value. Sci Signal. 2022;15(736):eabj9842.

62. Sood A, Lebel ME, Dong M, Fournier M, Vobecky SJ, Haddad E, et al. CD5 levels define functionally heterogeneous populations of naive human CD4(+) T cells. Eur J Immunol. 2021;51(6):1365–76.

63. Hogquist KA, and Jameson SC. The self-obsession of T cells: how TCR signaling thresholds affect fate ‘decisions’ and effector function. Nat Immunol. 2014;15(9):815–23.

64. Wadenpohl J, Seyfarth J, Hehenkamp P, Hoffmann M, Kummer S, Reinauer C, et al. CD5- expressing CD8(+) T-cell subsets differ between children with type 1 diabetes and controls. Immunol Cell Biol. 2021;99(10):1077–84.

65. Voisinne G, Gonzalez de Peredo A, and Roncagalli R. CD5, an Undercover Regulator of TCR Signaling. Front Immunol. 2018;9:2900.

66. Lozano F, Simarro M, Calvo J, Vila JM, Padilla O, Bowen MA, et al. CD5 signal transduction: positive or negative modulation of antigen receptor signaling. Crit Rev Immunol. 2000;20(4):347–58.

67. Matson CA, Choi S, Livak F, Zhao B, Mitra A, Love PE, et al. CD5 dynamically calibrates basal NF-kappaB signaling in T cells during thymic development and peripheral activation. Proc Natl Acad Sci U S A. 2020;117(25):14342–53.

68. Axtell RC, Xu L, Barnum SR, and Raman C. CD5-CK2 binding/activation-deficient mice are resistant to experimental autoimmune encephalomyelitis: protection is associated with diminished populations of IL-17-expressing T cells in the central nervous system. J Immunol. 2006;177(12):8542–9.

69. Wong CP, Li L, Frelinger JA, and Tisch R. Early autoimmune destruction of islet grafts is associated with a restricted repertoire of IGRP-specific CD8+ T cells in diabetic nonobese diabetic mice. J Immunol. 2006;176(3):1637–44.

70. Hong X, Meng S, Tang D, Wang T, Ding L, Yu H, et al. Single-Cell RNA Sequencing Reveals the Expansion of Cytotoxic CD4(+) T Lymphocytes and a Landscape of Immune Cells in Primary Sjogren’s Syndrome. Front Immunol. 2020;11:594658.

71. Rosati E, Pogorelyy MV, Dowds CM, Moller FT, Sorensen SB, Lebedev YB, et al. Identification of Disease-associated Traits and Clonotypes in the T Cell Receptor Repertoire of Monozygotic Twins Affected by Inflammatory Bowel Diseases. J Crohns Colitis. 2020;14(6):778–90.

72. Ihantola EL, Ilmonen H, Kailaanmaki A, Rytkonen-Nissinen M, Azam A, Maillere B, et al. Characterization of Proinsulin T Cell Epitopes Restricted by Type 1 Diabetes-Associated HLA Class II Molecules. J Immunol. 2020;204(9):2349–59.

73. Shizuru JA, Taylor-Edwards C, Livingstone A, and Fathman CG. Genetic dissection of T cell receptor V beta gene requirements for spontaneous murine diabetes. J Exp Med. 1991;174(3):633–8.

74. Wucherpfennig KW, and Gagnon E. Positively selecting peptides: their job does not end in the thymus. Nat Immunol. 2009;10(11):1143–4.

75. Daniels MA, Teixeiro E, Gill J, Hausmann B, Roubaty D, Holmberg K, et al. Thymic selection threshold defined by compartmentalization of Ras/MAPK signalling. Nature. 2006;444(7120):724–9.

76. Rogers D, Sood A, Wang H, van Beek JJP, Rademaker TJ, Artusa P, et al. Pre-existing chromatin accessibility and gene expression differences among naive CD4(+) T cells influence effector potential. Cell Rep. 2021;37(9):110064.

77. Drobek A, Moudra A, Mueller D, Huranova M, Horkova V, Pribikova M, et al. Strong homeostatic TCR signals induce formation of self-tolerant virtual memory CD8 T cells. EMBO J. 2018;37(14).

