## Supplementary material for "Thymic self-recognition-mediated TCR signal strength modulates antigen-specific CD8^+^ T cell pathogenicity in non-obese diabetic mice": V2_Supplementary material

**This PDF file includes:**

**Supplementary Figure 1.** Enhanced TCR signaling, transcription factor expression and cytokine profiles are unveiled in CD5^hi^CD8^+^ T cells of NOD mice.

**Supplementary Figure 2.** Gene expression patterns and enrichment are comparatively analyzed between naïve CD5^hi^ and CD5^lo^CD8^+^ T cells.

**Supplementary Figure 3.** Thymocyte development and respective selection profiles are compared between NOD and NOD8.3 mice.

**Supplementary Figure 4.** Thymocyte-stage self-reactivity correlates peripheral diabetogenic T cell proliferation profiles in NOD, BDC2.5 and NOD8.3 mice.

**Supplementary Figure 5.** Immunomodulatory effects of transgenic Pep on effector T cell functions are analyzed.

**Supplementary Figure 6.** The pairing of VJ gene segments in the TCR repertoire is compared between CD5^hi^CD8^+^ and CD5^lo^CD8^+^ cells.

**Supplementary Figure 7.** The genotyping results of NOD, dLPC/NOD, NOD8.3 and dLPC/NOD8.3 mice using the indicated F’ and R’ primers are shown.

**Supplementary Table 1.** 133 genes are upregulated in sorted naïve CD5^hi^CD8^+^ T cells compared to CD5^lo^CD8^+^ T cells.

**Supplementary Table 2.** 52 genes are downregulated in sorted naïve CD5^hi^CD8^+^ T cells compared to CD5^lo^CD8^+^ T cells.

**Supplementary Table 3.** Significantly enriched gene ontology terms in naïve CD5^hi^CD8^+^ compared to CD5^lo^CD8^+^ T cells are identified.

**Supplementary Table 4.** Reagent list.

**Supplementary Table 5.** The MiXCR pipeline executes the following steps for TCR repertoire analysis from RNA-seq data.

**Supplementary Table 6.** The MiXCR pipeline parameters and output paths using CD5hi_1_trimmed as an example are shown.

**Other supplementary material for this manuscript includes the following:**

**Supplementary data.** CD5hi_CD5lo_TCR_Repertoires_TRAV_TRBV_CDR3.xlsx

**Supplementary data.** CD5hi_1_trimmed.clonotypes.TRA.txt

**Supplementary data.** CD5hi_1_trimmed.clonotypes.TRB.txt

**Supplementary data.** CD5hi_2_trimmed.clonotypes.TRA.txt

**Supplementary data.** CD5hi_2_trimmed.clonotypes.TRB.txt

**Supplementary data.** CD5lo_1_trimmed.clonotypes.TRA.txt

**Supplementary data.** CD5lo_1_trimmed.clonotypes.TRB.txt

**Supplementary data.** CD5lo_2_trimmed.clonotypes.TRA.txt

**Supplementary data.** CD5lo_2_trimmed.clonotypes.TRB.txt

All of the above are available in the following GitHub repository: <https://github.com/Chia-Lo/TCR-signal-strength-modulates-antigen-specific-CD8-T-cell-pathogenicity-in-non-obese-diabetic-mice.git>

**Supplementary methods**

**Supplementary figure legends**

**Supplementary Figure 1. Enhanced TCR signaling, transcription factor expression and cytokine profiles are unveiled in CD5^hi^CD8^+^ T cells of NOD mice.** Representative FLOW analysis plots show a comparison of positive expression percentages in (**A**) proximal TCR signaling molecules p-CD3ζ and p-Erk, (**B**) transcription factors T-bet and Eomes, and (**C**) cytokines Granzyme B, TNF-α, IFN-γ and IL-2 within total CD8^+^ (grey), CD5^hi^CD8^+^ (red), CD5^lo^CD8^+^ (blue) cells, and fluorescence minus one (FMO) controls (light grey) from spleen (left) and PLNs (right).

**Supplementary methods**

*Assessment of insulitis and diabetes in NOD mice.* Pancreata were harvested from NOD mice with the indicated age in experiment, fixed in 10% neutral buffered formalin, and embedded in paraffin. Sections were cut and stained with hematoxylin and eosin. Insulitis was assessed by counting the number of islets showing infiltration of immune cells. Insulitis scoring criteria: 0, normal islet; 1, intact islet with few scattered mononuclear cells in the surroundings (<25% of the islet infiltrated); 2, peri-insulitis (25% to 50% of the islet infiltrated); 3, insulitis (50% to 75% of the islet infiltrated); 4, severe insulitis (75% to 100% of the islet infiltrated). Diabetes was assessed by monitoring urine glucose levels using a Chemstrips (Boehringer Mannheim, Indianapolis, IN). Mice with glucose levels above 500 mg/dL for two consecutive tests were considered diabetic.

*T cell proliferation assay.* To conduct proliferation assays, splenocytes or purified CD8^+^ T cells from 6-8-week-old normoglycemia female NOD mice were cultured in triplicate wells of 96-well flat-bottom plates (2–5 × 10^5^ cells/200 μl/well), along with indicated concentrations of soluble anti-CD3 (clone: 145-2C11, BD Pharmingen™), soluble anti-CD28 (clone: 37.51, BioLegend), or Dynabeads™ Mouse T-Activator CD3/CD28 (ThermoFisher, cat# 11456D). After the indicated incubation time, the cells were pulsed with 1 mCi [methyl-^3H] thymidine (PerkinElmer, Shelton, CT) per well and harvested following an additional 16–18 hours. The harvested samples were then transferred onto UniFilter-96, GF/C microplates (PerkinElmer), and the incorporated [methyl-^3H] thymidine was quantified using a Packard TopCount Microplate Scintillation Counter. For cell division assays, CD8^+^ T cells were isolated from pooled spleens and mLNs of 6-8-week-old normoglycemia female NOD mice using the Miltenyi Biotec MACS CD8a T Cell Isolation Kit. The cells were suspended in complete medium containing RPMI-1640, 10% FBS, 2 mM L-glutamine, 100 U/ml penicillin G, 0.1 mg/ml streptomycin and 10 mM HEPES, at a concentration of 1-2 million cells/ml. Cell labeling was performed using the fluorescent dye CellTrace™ Violet (CTV) as per the manufacturer's instructions (ThermoFisher, cat# C34557), followed by two washes with complete medium to remove excess CTV. Subsequently, cells were plated in 96-well plates coated with anti-CD3 and anti-CD28 antibodies at various concentrations as indicated in the figure, or in uncoated wells as a negative control. The cells were then incubated for 2 days at 37°C, 5% CO2, in the presence of IL-2, IL-7 and IL-15 cytokines (10 ng/ml each). Following the incubation period, cells were harvested and analyzed using flow cytometry to evaluate proliferation based on CTV dilution and determine the impact of different levels of anti-CD3 and anti-CD28 stimulation on T cell proliferation. Dead cells were excluded from the analysis using a viability stain (Zombie Red™, BioLegend).

*Cytokine staining*. For intracellular cytokine detection, splenocytes and PLN-derived T cells were stimulated *ex vivo* with ionomycin (1 μg/ml) and PMA (50 ng/ml) in the presence of monensin (2 μM) for 4 hours at 37°C. Following stimulation, cells were first stained for surface markers, fixed, and permeabilized using the Fixation/Permeabilization kit (eBioscience). Intracellular cytokines were detected using the following fluorochrome-conjugated antibodies: IFN-γ (XMG1.2), granzyme B (NGZB), TNF-α (MP6-XT22) and IL-2 (JES6-5H4). Data were acquired using Attune NxT V4 flow cytometer and analyzed with FlowJo software. Unstimulated controls were included to assess background cytokine expression.

*Flow cytometric analysis of immune cell markers.* For flow cytometric analysis of immune cell markers, single-cell suspensions were prepared from spleens and lymph nodes. The cells were then stained with a panel of surface antibodies, followed by intracellular staining. Analysis of the stained cells was performed using an Attune NxT V4 flow cytometer and FlowJo software (v10.4.0). The panel of antibodies used for flow cytometric analysis included the following markers: CD3 (17A2), CD4 (RM4-5), CD8a (53-6.7), CD5 (53-7.3), CD25 (PC61.5), CD44 (IM7), CD62L (MEL-14), CD69 (H1.2F3), CD122 (TM-β1), CD127 (A7R34), CXCR3 (CXCR3-173) and PD-1 (29F.1A12) for surface markers. Intracellular markers included Foxp3 (FJK-16s), IFN-γ (XMG1.2), ERK1/2 (pT202/pY204), Eomes (Dan11mag), T-bet (eBio4B10), granzyme B (NGZB), TNF-α (MP6-XT22), IL-2 (JES6-5H4) and p-CD247 (H4B4) (Supplementary Table 4).

*Transcriptomic analysis of CD5^hi^CD8^+^ and CD5^lo^CD8^+^ T cell subpopulations in NOD mice.* Single-cell suspensions were prepared from the spleens and lymph nodes of NOD mice. The cells were then pre-purified using the CD8a (CD8^+^) T Cell Isolation Kit from Miltenyi Biotec (cat# 130-104-075), followed by sorting of CD5^hi^CD8^+^ and CD5^lo^CD8^+^ T cells using a FACS Aria II cell sorter. The sorted cells were then centrifuged, and the resulting cell pellets were stored at -80°C for subsequent RNA extraction. RNA extraction was performed using the Qiagen RNeasy Plus Micro Kit, and RNA quality was evaluated using an Agilent 2100 Bioanalyzer. Subsequently, cDNA libraries were generated using the NEBNext Ultra II RNA Library Prep Kit, and sequencing was carried out on an Illumina HiSeq 2500 platform. The generated sequencing reads underwent processing and analysis using bioinformatics tools, including STAR, HTSeq, and DESeq2. Differential gene expression analysis was performed to identify genes exhibiting significant expression changes between CD5^hi^CD8^+^ and CD5^lo^CD8^+^ T cells. Additionally, KEGG pathway analysis was conducted to identify enriched metabolic biological pathways associated with these differential gene expression patterns.

*Isolation of pancreas-infiltrating cells.* The pancreata were excised from NOD mice and placed in a tube containing RPMI 1640 medium without FBS, supplemented with 1 mg/mL collagenase XI (Sigma-Aldrich). Subsequently, the pancreas tissue was mechanically dissociated using a scalpel to obtain a single-cell suspension. The dissociated tissue was then incubated for 30-40 minutes at 37°C with intermittent agitation to allow for enzymatic digestion. Following digestion, the contents were filtered through a cell strainer to remove debris and large aggregates. A density gradient centrifugation using Histopaque 1077 (Sigma-Aldrich) was performed to isolate the pancreas-infiltrating cells. The interphase containing the pancreas-infiltrating cells was carefully collected, and the cells were incubated with Cell Dissociation Buffer (Life Technologies, catalog number 41966-029) and washed again with RPMI 1640 medium. Subsequently, the resulting cells were counted and prepared for downstream flow cytometry analysis.

*Adoptive transfer of sorted CD5^hi^CD8^+^ and CD5^lo^CD8^+^ T cells.* Naive CD8^+^ T cells with 10% CD5 upper and lower distribution were sorted from spleens and lymph nodes of NOD mice using a FACS Aria II cell sorter. The sorted two groups (1x10^6^ cells for each) were then respectively combined with 2x10^6^ CD4^+^ T cells from other NOD mice. The sorted cells were then centrifuged and resuspended in PBS. NOD Rag1^-/-^ recipients were transferred with each sorted population by intraperitoneal (IP) injection. After 4 weeks transfer, the spleens and lymph nodes of recipient mice were harvested and analyzed for the presence of transferred cells using flow cytometry, or NOD Rag1^-/-^ recipients were monitored for diabetes onset by measuring urine glucose levels twice a week using a Chemstrips. Diabetes was defined as urine glucose levels above 500 mg/dl on two consecutive measurements.

*TCR repertoire analysis by MiXCR pipeline on bulk RNA sequencing data.* RNA extraction, cDNA library preparation, sequencing, and data preprocessing following the manufacturer's instructions were described as previously in bulk RNA seq for DEGs analysis. The obtained preprocessed RNA sequencing data were then analyzed using the MiXCR pipeline (3.0.8) (77). The following steps outline the upstream process: First, RNA-seq reads were aligned to the reference genome using the MiXCR aligner with the specified parameters, including allowing partial alignments and specifying the species and library. After alignment, refinement was performed to enhance the quality of the alignments. Subsequently, partial assembly of the aligned reads was carried out to identify TCR sequences, followed by extension of the CDR3 of the TCR sequences for improved accuracy. Clonotypes were then assembled based on the extended CDR3 sequences to identify unique TCR clones, and contig assembly was performed to reconstruct full-length TCR sequences from the assembled clonotypes. Notably, intermediate files generated during the upstream analysis steps were not available due to file management constraints (Supplementary Table 5). Finally, the assembled TCR repertoire data from MiXCR including T cell receptor α (TRA) and β (TRB) variable gene sequences were exported by online VDJviz for downstream analysis (78) (CD5hi_CD5lo_TCR_Repertoires_TRAV_TRBV_CDR3.xlsx, available at <https://github.com/Chia-Lo/TCR-signal-strength-modulates-antigen-specific-CD8-T-cell-pathogenicity-in-non-obese-diabetic-mice.git>). The output of the MiXCR analysis provided information on the clonotype sequences and their respective frequencies for each TCR clonotype. Specific output paths for samples, including OutputPath, ForwardReadFile, and ReverseReadFile, are detailed in the MiXCR pipeline parameters section using the CD5hi_1_trimmed sample as an example (Supplementary Table 6).

*TCR repertoire visualization by VDJviz and Immunarch.* The output files from MiXCR analysis were utilized for subsequent visualization. The clonotype files were imported into VDJviz (1.0.3.2) (78). VDJviz was used to explore key repertoire characteristics, including spectratype profiles, V-(D)-J recombination patterns and top 20 CDR3α and CDR3β motifs between CD5^hi^ and CD5^lo^ populations. Additionally, the exported TRA or TRB files from the MiXCR pipeline or VDJviz analysis were processed using R packages rTCRBCRr (1.0.0), readr (2.1.5), ggplot2 (3.4.4), dplyr (1.1.4), tidyr (1.3.1), readxl (1.4.3) and writexl (1.4.2). These packages were utilized for visualizing V and J gene usage in the TCR α and β chain, analyzing CDR3 length distribution and repertoire overlap, and examining CDR3 biochemical properties and diversity between CD5^hi^ and CD5^lo^ T cell populations.

*Hydrophobicity analysis of CDR3 amino acids*. In this study, the 20 naturally occurring amino acids were clustered into three subgroups based on their hydropathic properties: hydrophobic (including "A," "C," "F," "I," "L," "M," "W," and "V"), neutral (including "G," "H," "P," "S," "T," and "Y"), and hydrophilic (including "E," "D," "K," "N," "Q," and "R"). These groupings were determined following the Kyte–Doolittle hydropathy scale (79). To calculate the hydrophobicity score of CDR3 amino acids, we utilized a custom-built function called calculateHydrophobicity, which is available at <https://github.com/Chia-Lo/TCR-signal-strength-modulates-antigen-specific-CD8-T-cell-pathogenicity-in-non-obese-diabetic-mice.git>. This function utilized the hydropathy scale dictionary to evaluate the hydrophobicity of each amino acid in the CDR3 sequences of CD5^hi^CD8^+^ and CD5^lo^CD8^+^ T cell populations. The calculated average hydrophobicity scores for each sample were then analyzed to discern differences in hydropathic properties between the two T cell subsets. Finally, the results were visualized using appropriate statistical methods to facilitate a comparative analysis of hydrophobicity scores between CD5^hi^CD8^+^ and CD5^lo^CD8^+^ T cells.

*Clonotype repertoire metrics*. Clonotype repertoire diversity was calculated using Shannon entropy, which accounts for both abundance and distribution of unique clonotypes in a sample. Shannon entropy was computed using the formula: -$\sum_{i=1}^{N} pi log(pi)$, where *pi* represents the frequency of each clonotype, and *N* is the number of unique clonotypes. These calculations were implemented using the calculate_repertoire_metrics function from the R package rTCRBCRr, which returns a vector containing diversity values based on a named vector of clonotype frequencies.

*Construction of the overlapped count matrix*. To evaluate the overlap of TCR clonotypes between CD5^hi^CD8^+^ and CD5^lo^CD8^+^ T cell populations, TCR sequences were obtained from sorted cells, and clonotypes were defined based on unique combinations of V gene, J gene, and CDR3 nucleotide sequences. For each sample, clonotype frequency was calculated by dividing the number of reads corresponding to each clonotype by the total TCR reads in that sample. An Overlapped Count Matrix was then generated, with clonotypes as rows and samples as columns, where each entry represented the frequency of a given clonotype in a specific sample, and undetected clonotypes were assigned a frequency of zero. The matrix was visualized as a heatmap to illustrate the repertoire overlap and distinguish shared versus unique clonotypes between CD5^hi^ and CD5^lo^ subsets. The R code used to generate the Overlapped Count Matrix is available at: <https://github.com/Chia-Lo/TCR-signal-strength-modulates-antigen-specific-CD8-T-cell-pathogenicity-in-non-obese-diabetic-mice.git>.
